# miR-940 suppresses ferroptosis by controlling expression of key regulatory genes

**DOI:** 10.64898/2026.02.09.704912

**Authors:** Andrea Kolak, Juliane Tschuck, Stefanie A. I. Weiß, Daniel Kaemena, Karolin Klimm, Ana Galhoz, Larissa Ringelstetter, Myles Fennell, Juliane Merl-Pham, Anna Artati, Stefanie Strasser, Ralph Garippa, Michael Witting, Hans Zischka, Joel A. Schick, Stefanie M. Hauck, Michael P. Menden, Michelle Vincendeau, Brent R. Stockwell, Kamyar Hadian

## Abstract

Ferroptosis is a form of regulated cell death that is characterized by iron-dependent lipid peroxidation. This process is regulated by specific metabolites, the lipid composition of the cells, redox-active iron, and antioxidant mechanisms. Although numerous regulators have been identified over the past decade, exploring other mechanisms, particularly from non-coding genomic regions, can build a thorough understanding of the multifaceted regulatory processes underlying ferroptosis. MicroRNAs (miRNAs) play a crucial role in gene regulation and cellular functions. Through a CRISPR KO screen, we identified miR-940 as a negative regulator of ferroptosis. Overexpression of miR-940 in several cell lines consistently suppressed ferroptosis induced by system x_c_^-^ inhibition. Notably, multiple cancer patient cohorts with elevated miR-940 levels exhibit reduced survival. Integrated bioinformatic, transcriptomic, and proteomic analyses revealed that miR-940 decreases the expression of ACSL4, LPCAT3, DMT1, and NCOA4, and simultaneously increases levels of GPX4. Pharmacological inhibition of GPX4 attenuated the protective effect of miR-940, indicating that its primary anti-ferroptotic activity is mediated through GPX4. Overall, this gene rewiring is associated with reduced levels of redox-active iron and diminished lipid peroxidation, consistent with ferroptosis suppression. These findings suggest that miR-940 coordinates ferroptosis inhibition, which presents a novel regulatory layer for therapeutic exploration in susceptible cancers.

## 1. Introduction

Cell homeostasis is dependent on cell division and cell death. The past two decades have shown that besides apoptosis, several other cell death modalities are executed in a regulated fashion [1]. Ferroptosis is a regulated cell death pathway that is based on iron-dependent lipid peroxidation [2-4]. It is initiated by the presence of phospholipids with polyunsaturated fatty acyl tails (PUFA-PLs), redox-active iron, and a defective lipid peroxide repair system [5]. PUFAs are incorporated into phospholipids by the action of the enzymes acyl-CoA synthetase long-chain family member 4 (ACSL4) and lysophosphatidylcholine acyltransferase 3 (LPCAT3) [6]. Iron is transported into cells by endocytosis mediated by transferrin receptor protein 1 (TfR1) and released from endosomes by divalent metal transporter 1 (DMT1) [4]. Iron is then stored in a non-redox-active form in iron-ferritin complexes and released from ferritin in a process called ferritinophagy, mediated by nuclear receptor coactivator 4 (NCOA4) [6]. Interestingly, TfR1 has also been shown to act as a potent marker to detect ferroptosis [7-8]. To prevent ferroptosis, cells have developed several ways to protect themselves from lipid peroxidation. The system x_c_^-^/GPX4/glutathione axis is the central ferroptosis inhibitory mechanism, where glutathione peroxidase 4 (GPX4) uses glutathione (GSH) to reduce lipid hydroperoxides to the corresponding alcohol forms [4]. In addition, there are at least two ferroptosis inhibitory modules that act independently of GPX4, namely the FSP1/ubiquinol/vitamin-K axis [9-11], and the GCH1/DHFR/tetrahydrobiopterin axis [12-13]. Several synthetic ferroptosis inducers (IKE, RSL3, ML162 and FINO_2_) as well as inhibitors (Fer-1 and Lip-1) have been developed that can modulate ferroptosis sensitivity [1]. Additionally, approved drugs have been repurposed to inhibit ferroptosis, such as seratrodast [14].

Several transcriptional factors control the expression of key ferroptosis regulators to balance ferroptosis induction. Examples are the nuclear factor-erythroid 2-related factor 2 (NRF2) [6], as well as nuclear receptors including the farnesoid X receptor (FXR) [15], the retinoic acid receptor (RAR) [16] or the estrogen receptor (ER) [17-18]. Gene expression can also be regulated by microRNAs (miRNAs), which either degrade the mRNA or target the untranslated regions (UTRs) of mRNA transcripts to prevent their translation [19]. Recent years have seen growing interest in miRNA-mediated regulation in many fields of research including ferroptosis. Although several studies have identified specific miRNAs that influence the ability of cells to undergo ferroptosis [20], there is an opportunity to expand our knowledge of this class of regulators.

The microRNA miR-940 is located on chromosome 16p13.3 and is known to exhibit dysregulated expression in various diseases, including many types of cancer [21]. It binds to the 3’ untranslated region (UTR) of target protein-coding genes, long non-coding RNAs, and circular RNAs, thereby affecting their transcription or post-transcriptional regulation [21]. Unlike many existing protein regulators that control tumor progression, the expression of miR-940 can be increased or decreased in cancer, depending on the tissue and cell type. Its expression is decreased in hepatocellular carcinoma [22] and non-small-cell lung cancer [23], but increased in gastric [24] and cervical cancer [25]. This opposing role of miR-940 has also been described in tumor metastasis [26], cell cycle progression [25], the epithelial-mesenchymal transition [27], among others. MiR-940 affects signaling pathways by either directly targeting molecules such as MAPK1 [28], or indirectly by inhibiting CBL-b and c-CBL, resulting in PD-L1 upregulation and an increase in STAT3 and AKT [29]. In addition to signal transduction, potential mechanisms by which miR-940 regulates tumorigenesis include the induction of DNA damage [30], interference with the folate metabolism through the downregulation of MTHFD2 [31], and the silencing of transcription factors such as ZNF24 [24]. However, a connection to ferroptosis regulation has never been reported.

In this study, we identified miR-940 as a key modulator of ferroptosis by systematically integrating data from a CRISPR KO screen, transcriptomics and proteomics. We demonstrate that miR-940 upregulates GPX4, while concurrently reducing levels of ferroptosis-promoting genes including ACSL4, LPCAT3, DMT1, and NCOA4. Together, this regulation by miR-940 reduces ferroptosis.

## 2. Results

### 2.1. CRISPR KO screen identifies miR-940 as a negative regulator of ferroptosis

To discover novel ferroptosis regulators, we performed a genome-wide CRISPR knockout (KO) screen in this study using the GeCKO v2 human CRISPR knockout pooled library [32]. Cas9-expressing HT-1080 cells, a widely used ferroptosis model, were transduced with the library and treated with the ferroptosis inducer Imidazole ketone erastin (IKE) [33] or DMSO. After 18 h, genomic DNA of surviving cells was extracted and subjected to next generation sequencing (NGS) (Figure 1A). Analysis of the abundance of gRNAs associated with miRNAs revealed a set of 30 differentially expressed gRNAs upon ferroptosis induction. Here, 15 of these were significantly enriched, suggesting that the corresponding miRNAs are pro-ferroptotic, and the other 15 were significantly depleted, indicating that the corresponding miRNAs are negative regulators of ferroptosis (Figure 1B).

**Figure 1.**
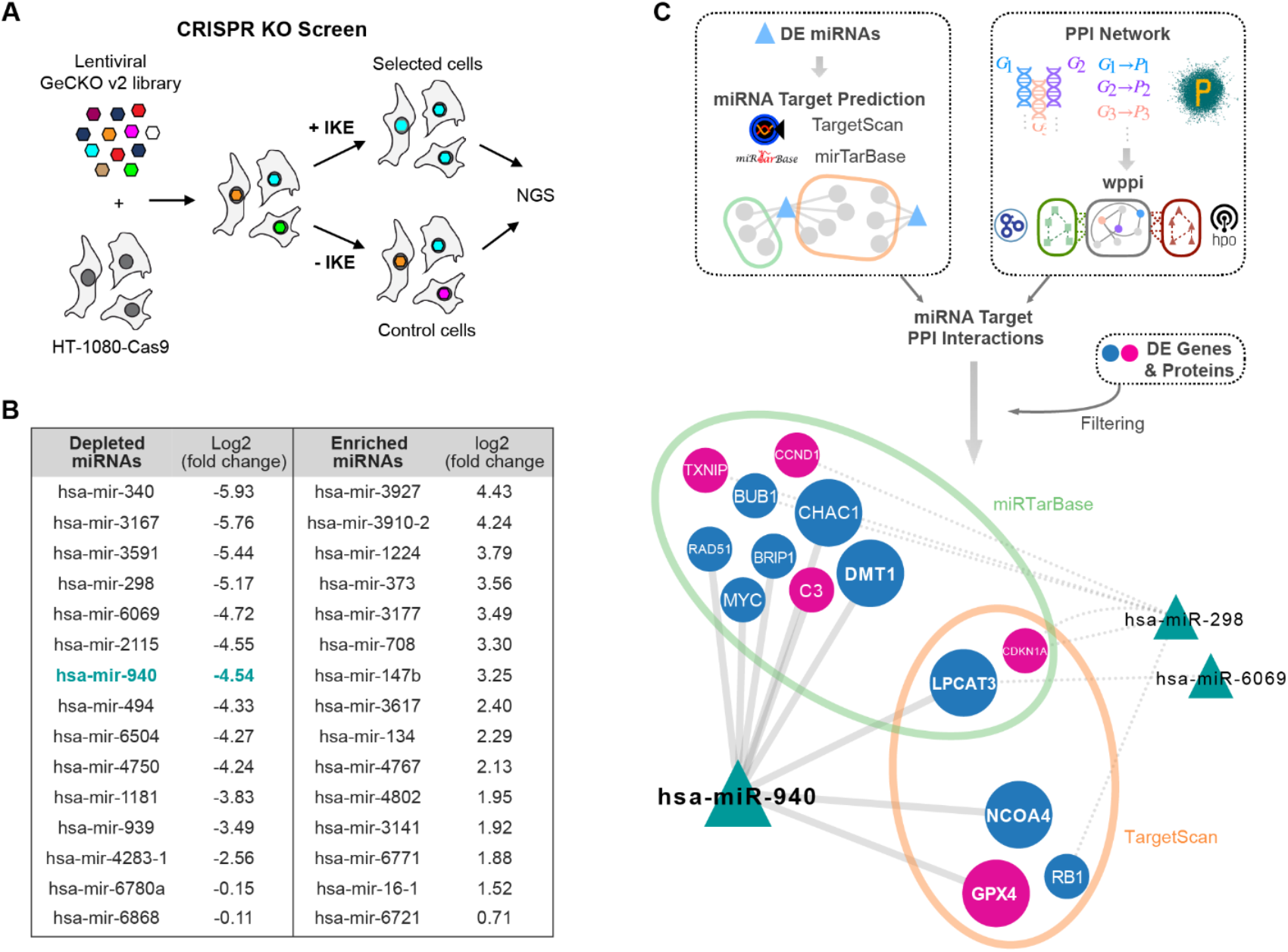
CRISPR KO screen identifies miR-940 as a negative regulator of ferroptosis. A) Illustration of the genome-wide CRISPR knockout screen using the GeCKO v2 human CRISPR knockout pooled library. B) Differentially regulated miRNAs resulting from the CRISPR knockout screen. C) Illustration of the bioinformatic pipeline for miRNA target prediction and protein-protein interaction (PPI) analysis. Gene targets of the differentially regulated miRNAs were retrieved from the TargetScan and miRTarBase databases. Validated gene targets from miRTarBase were leveraged in a weighted PPI (WPPI) network analysis to expand the candidate gene list. Focusing on targets among differentially expressed genes and proteins (FDR = 1% and |log2 fold-change| > 1), a total of 14 genes were identified as targets of hsa-miR-940, hsa-miR-298 and hsa-miR-6069. In the network illustration, blue nodes indicate up-regulated genes, red nodes indicate down-regulated genes and triangles represent miRNAs. Large nodes represent the 5 genes identified by miRTarBase and TargetScan (i.e., CHAC1, DMT1, LPCAT3, NCOA4 and GPX4), while small nodes represent the 9 new genes identified through the WPPI network analysis (i.e., TXNIP, CCND1, BUB1, RAD51, BRIP1, MYC, C3, CDKN1A and RB1).

To predict most promising miRNA candidates that may be involved in ferroptosis-related pathways, we performed an in-depth bioinformatics analysis. We seeded 95 genes/proteins with a known contribution in ferroptosis regulation into a functional network analysis (Figure 1C). We then performed miRNA target prediction and protein-protein interaction (PPI) analysis, ranking the candidate genes based on their association score with the ferroptosis-related seed genes. These analyses revealed that miR-940 showed the strongest association with ferroptosis regulatory proteins including DMT1, LPCAT3, NCOA4 and GPX4 (Figure 1C). This prompted us to select this miRNA for further analysis.

### 2.2. miR-940 inhibits IKE-induced ferroptosis

Previous studies suggested that miR-940 may induce apoptosis [34-35]. In contrast, using Annexin V/7-AAD staining, we show that miR-940 does not induce apoptosis in our experimental setting (Figure 2A). Next, we performed a series of experiments to confirm a negative regulatory role of miR-940 in ferroptosis. We transfected HT-1080, HepG2 cells and HEK293T cells with the precursor miRNA of miR-940 (pre-miR-940) and detected a robust upregulation of miR-940 transcripts (Supplementary Figure 1A). We used three different cell lines from three different tissue origins to demonstrate the generalizability of ferroptosis suppression by miR-940. We then induced ferroptosis using IKE, RSL3, ML162 or FINO_2_. These molecules all selectively induced ferroptosis, because the specific ferroptosis inhibitor Fer-1 fully restored cell viability when co-treated with them (Supplementary Figure 1B). As expected from the screening results, ferroptosis induction by IKE was significantly rescued by miR-940 overexpression in HT-1080 cells (Figure 2B). Interestingly, ferroptosis induced by GPX4 inhibition (RSL3, ML162 or FINO_2_) was only slightly inhibited by miR-940 (Figure 2B). IKE-induced ferroptosis inhibition by miR-940 was additionally demonstrated through PI staining (Figure 2C) and crystal violet staining (Figure 2D). Importantly, staurosporine-induced apoptosis was not inhibited by miR-940 in HT-1080 cells (Figure 2E), suggesting specificity towards ferroptosis over apoptosis. These data could be confirmed using Annexin V/7-AAD staining (Supplementary Figure 1C). To determine if this effect was cell line-specific, we examined ferroptosis susceptibility after miR-940 overexpression in two additional cell lines. HepG2 cells were also rescued by miR-940 from IKE-induced ferroptosis, but not after RSL3, ML162, and FINO_2_ treatment (Figure 2F). Similar results were obtained in HEK293T cells, namely, suppression of ferroptosis by miR-940 upon IKE-induced system x_c_^-^ inhibition only (Figure 2G). In both cell lines, HepG2 and HEK293T cells, miR-940 had no effect on apoptosis (Figure 2H-I).

**Figure 2.**
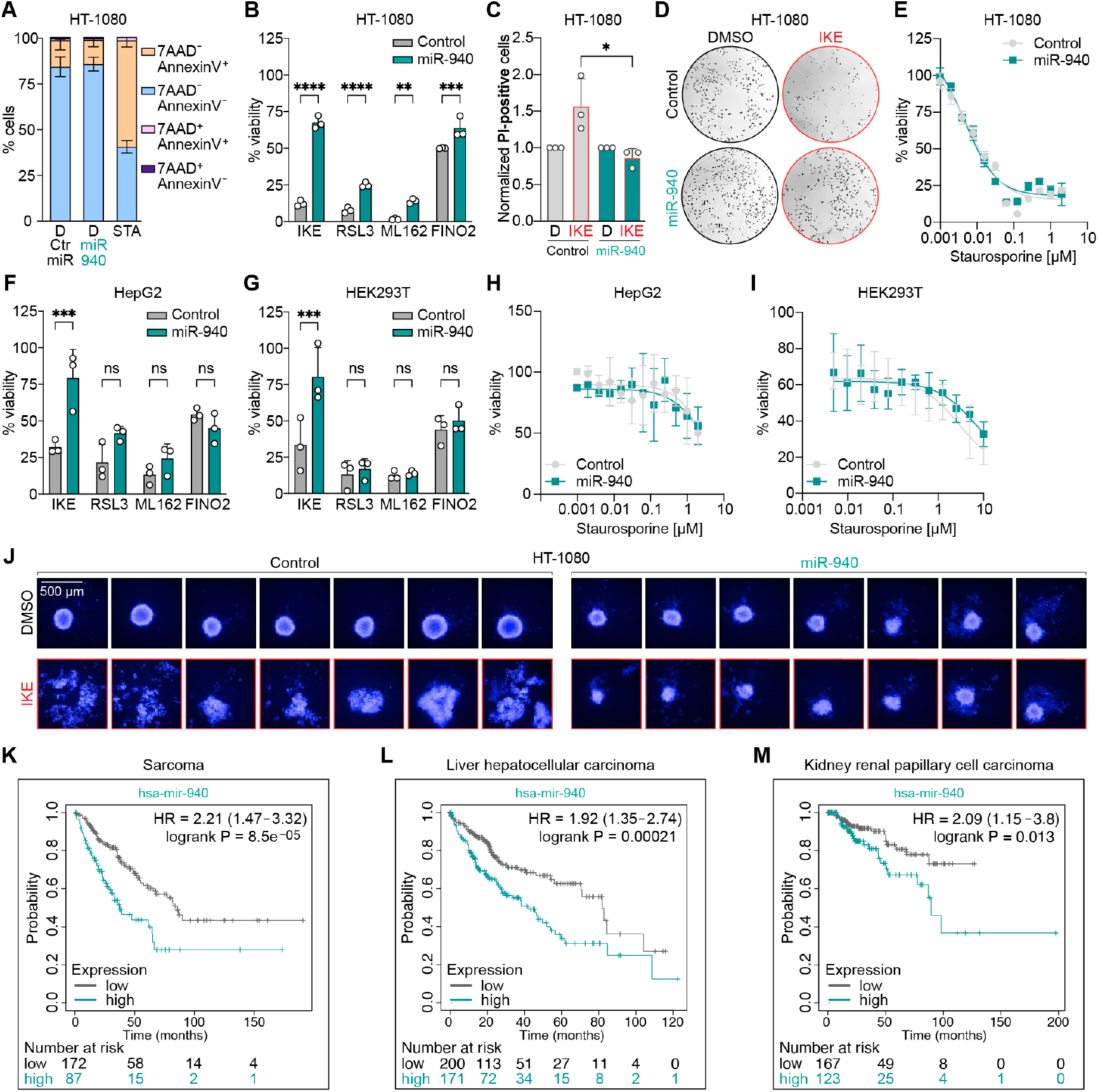
miR-940 inhibits IKE-induced ferroptosis. A) AnnexinV/7-AAD staining in pre-miR-940- and control-transfected HT-1080 cells 24 h post transfection. AnnexinV positive control was included by the addition of staurosporine (STA, 1 µM), data are mean ± SD of n = 3 biological replicates. B) Inhibition of ferroptosis in pre-miR-940- and control-transfected HT-1080 cells treated with 0.3 µM IKE, 0.25 µM RSL3, 0.25 µM ML162 and 0.6 µM FINO_2_; **** = p ≤ 0.0001, *** = p ≤ 0.001, ** = p ≤ 0.01, 2way ANOVA, data are mean ± SD of n = 3 biological replicates. C) Measurement of PI-positive cells upon induction of ferroptosis (2 µM IKE) in pre-miR-940- and control-transfected HT-1080 cells, * = p ≤ 0.05, 2way ANOVA, data are mean ± SD of n = 3 biological replicates. D) Crystal violet staining of pre-miR-940- and control-transfected HT-1080 cells after treatment with 2 µM IKE for 24 h. E) Dose dependent induction of apoptosis with staurosporine in pre-miR-940- and control-transfected HT-1080 cells. F,G) Inhibition of ferroptosis in pre-miR-940- and control-transfected HepG2 (F) and HEK293T (G) cells treated with (F) 2.5 µM IKE, 0.5 µM RSL3, 0.25 µM ML162 and 2.5 µM FINO_2_ and (G) 0.6 µM IKE, 0.25 µM RSL3, 0.25 µM ML162 and 1.25 µM FINO_2_; *** = p ≤ 0.001, ns = p > 0.05, 2way ANOVA, data are mean ± SD of n = 3 biological replicates. H,I) Dose dependent induction of apoptosis with staurosporine in pre-miR-940- and control-transfected HepG2 (H) and HEK293T (I) cells. J) 3D spheroid formation of pre-miR-940- and control-transfected HT-1080 cells treated with 2 µM IKE, median intensity of Hoechst signal visualized. K,L,M) Query of Kaplan-Meier (KM) plotter to analyze the correlation of miR-940 expression and patient survival in sarcoma (K), liver hepatocellular carcinoma (L) and kidney renal papillary cell carcinoma (M).

We next tested the inhibitory effect of miR-940 on ferroptosis in a more physiological setting. Therefore, we used 3D spheroids of HT-1080 cells, either transfected with pre-miR-940 or the control (seven spheroids per condition are depicted in Figure 2J). Spheroid formation in cells treated with IKE was impaired compared to the DMSO control. This effect could not be observed in cells transfected with pre-miR-940, which showed no difference in the spheroid integrity and size between DMSO and IKE treatment (Figure 2J). Notably, more cells migrated out of the spheroids when transfected with pre-miR-940, which is likely the result of altered pathways by miR-940, thereby facilitating cell migration [36]. Hence, we validated the anti-ferroptotic effect of miR-940 also in a three-dimensional setup that more closely reflects physiological conditions. We also queried the KM Plotter [37-38] and analyzed the correlation between miR-940 expression and patient survival. Patients with higher levels had worse survival rates in sarcoma (Figure 2K), liver hepatocellular carcinoma (Figure 2L), and kidney renal papillary cell carcinoma (Figure 2M). These patient data align well with our HT-1080 (fibrosarcoma), HepG2 cells (liver hepatocellular carcinoma) and HEK293T (kidney cells), which exhibit ferroptosis suppression by miR-940.

### miR-940 modulates key regulators of ferroptotic cell death

To further investigate the role of miR-940 in ferroptosis-related pathways, we subjected total RNA from pre-miR-940- and control-transfected cells to RNA sequencing (RNAseq). Transcriptomics analysis revealed the downregulation of four ferroptosis-promoting target genes: LPCAT3, DMT1, NCOA4, and ACSL4 (Figure 3A). This largely aligned with the prediction of our bioinformatics pipeline (Figure 1C). Transcriptomics also showed that GPX4, the central gatekeeper of ferroptosis, was upregulated upon pre-miR-940 overexpression (Figure 3A), which aligns with the bioinformatic prediction. Normalized transcript counts from the RNAseq data are shown in Figure 3B. We did not assess direct 3’ UTR binding for these transcripts; thus, direct miR-940-mRNA interactions remain to be established. However, predicted miR-940 binding sites were identified in the 3′ UTRs of the ferroptosis regulators *NCOA4, ACSL4, LPCAT3, DMT1*, and *GPX4* (Supplementary Fig. 2 and Supplementary Table 1). Notably, *NCOA4* revealed a high-confidence 8mer site predicted by both TargetScan and miRDB, while *DMT1* was previously validated as a direct miR-940 target [39]. Along with transcriptomics, we performed proteomics and observed downregulation of ACSL4 protein levels. Unfortunately, the other proteins were not detected due to limited resolution (Supplementary Fig. 3).

**Figure 3.**
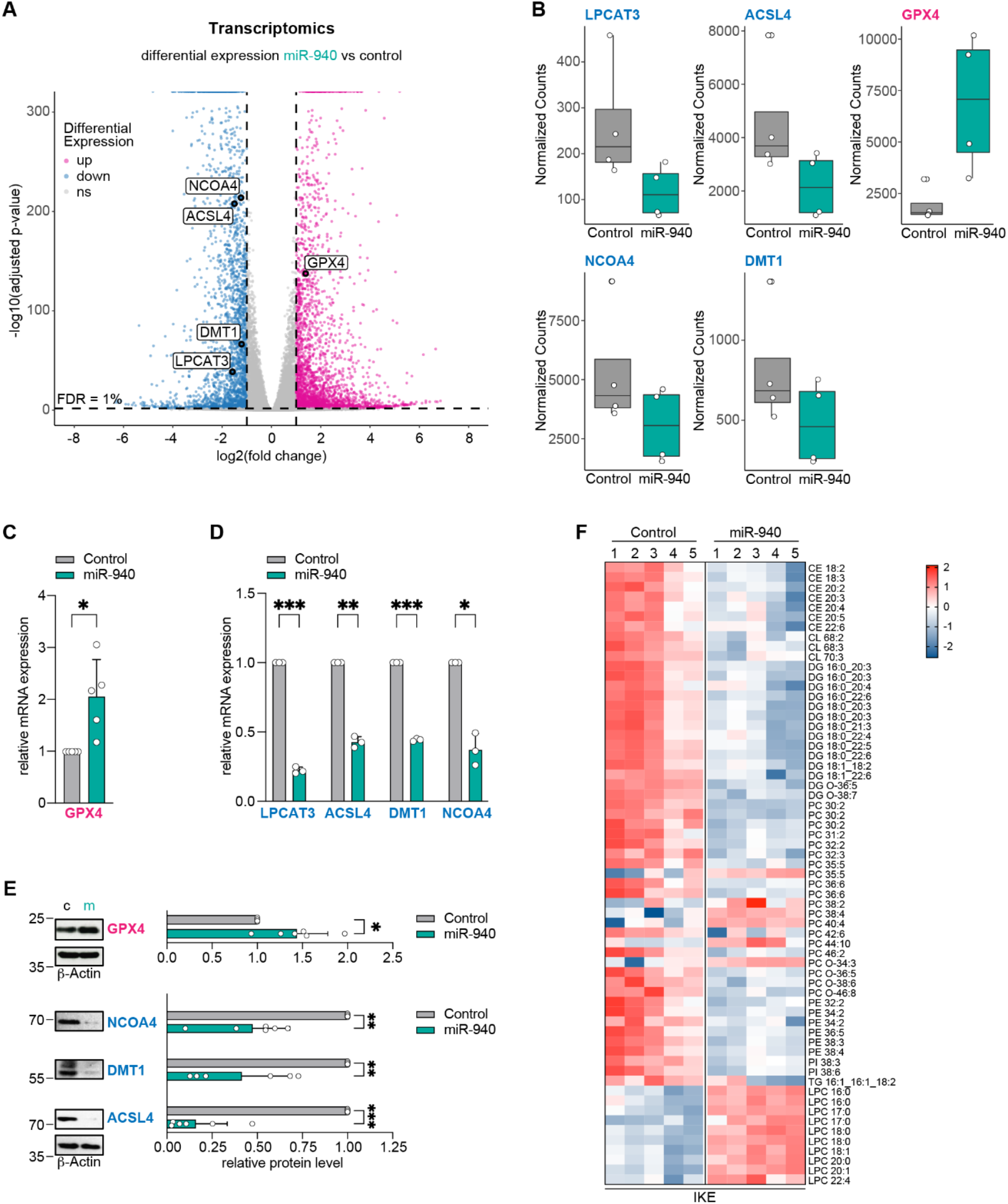
miR-940 modulates key regulators of ferroptotic cell death. A) Transcriptomics analysis of miR-940-overexpressing HT-1080 cells show downregulation of NCOA4, ACSL4, DMT1 and LPCAT3 and upregulation of GPX4, significance at False Discovery Rate (FDR) of 1% and absolute value of log2 fold-change (log2FC) > 1, Wald test adjusted with Benjamini-Hochberg correction. B) Normalized counts of transcriptomics analysis of LPCAT3, ACSL4, DMT1, NCOA4, and GPX4, data are mean ± SD of n = 4 biological replicates. C) qRT-PCR results of GPX4 in miR-940 transfected cells, * = p ≤ 0.05, unpaired t-test with Welch’s correction, data are mean ± SD of n = 3 biological replicates. C) qRT-PCR results of GPX4 in miR-940 transfected cells, * = p ≤ 0.05, unpaired t-test with Welch’s correction, data are mean ± SD of n = 5 biological replicates. D) qRT-PCR results of LPCAT3, ACSL4, DMT1 and NCOA4 in miR-940-overexpressing cells, * = p ≤ 0.05, ** = p ≤ 0.01, *** = p ≤ 0.001, unpaired t-test with Welch correction, data are mean ± SD of n = 3 biological replicates. E) Western Blot analysis of miR-940-overexpressing HT-1080 cells with staining against GPX4, NCOA4, DMT1, and ACSL4 relative to β-Actin levels, * = p ≤ 0.05, ** = p ≤ 0.01, *** = p ≤ 0.001, unpaired t-test with Welch correction, data are mean ± SD of n = 6 biological replicates. F) Lipidomics analyses of pre-miR-940- and control-transfected HT-1080 cells treated with 2 µM IKE show depletion of ferroptosis-relevant PUFA-containing phospholipids upon miR-940 overexpression, CE cholesteryl ester, CL cardiolipin, DG diacylglyceride, PC phosphatidylcholine, PE phosphatidylethanolamine, PI phosphatidylinositol, TG triglyceride, LPC lysophosphatidylcholin, depicted are Z-scores with p ≤ 0.05, two-tailed t-test, n = 5 biological replicates.

To confirm the omics data, the mRNA levels of all differentially expressed target regulators were analyzed by qRT-PCR. Significant upregulation of GPX4 (Figure 3C) and downregulation of LPCAT3, ACSL4, DMT1 and NCOA4 (Figure 3D) could be confirmed upon transfection with pre-miR-940. Similar results were obtained in HepG2 cells and HEK293T cells (Supplementary Fig. 4A-B). Finally, we investigated whether miR-940 reduces the protein levels of the aforementioned target proteins by performing western blot analysis and quantification. Overexpression of miR-940 significantly increased GPX4 protein levels, protein levels of NCOA4, DMT1 and ACSL4 were significantly decreased (Figure 3E). These findings demonstrate that miR-940 modulates key regulators of ferroptotic cell death, as detected on RNA as well as protein levels. Interestingly, lipidomics analyses of control-miRNA-versus miR-940-transfected cells demonstrate that several ferroptosis-relevant PUFA-containing phospholipids are depleted upon miR-940 overexpression (Figure 3F). According to our data, this largely correlates with reduction in ACSL4 and LPCAT3 levels. Yet, reflecting on the data that miR-940 largely failed to rescue from ferroptosis upon GPX4 inhibition, this fact sets GPX4 as the major mechanism by which miR-940 suppresses ferroptosis, because upregulated GPX4 proteins (Figure 3E) are still inhibited by RSL3, ML162 and FINO_2_.

### 2.3. miR-940 suppresses iron accumulation and lipid peroxidation

To assess the mechanism behind the anti-ferroptotic actions of miR-940, we first investigated glutathione levels within HT-1080 cells following treatment with IKE. No significant increase in GSH concentration was observed in miR-940-transfected versus control cells. Both, miR-940-transfected cells and control cells, showed decreased GSH levels upon IKE-induced ferroptosis, indicating that miR-940 has no direct influence on GSH levels (Figure 4A).

**Figure 4.**
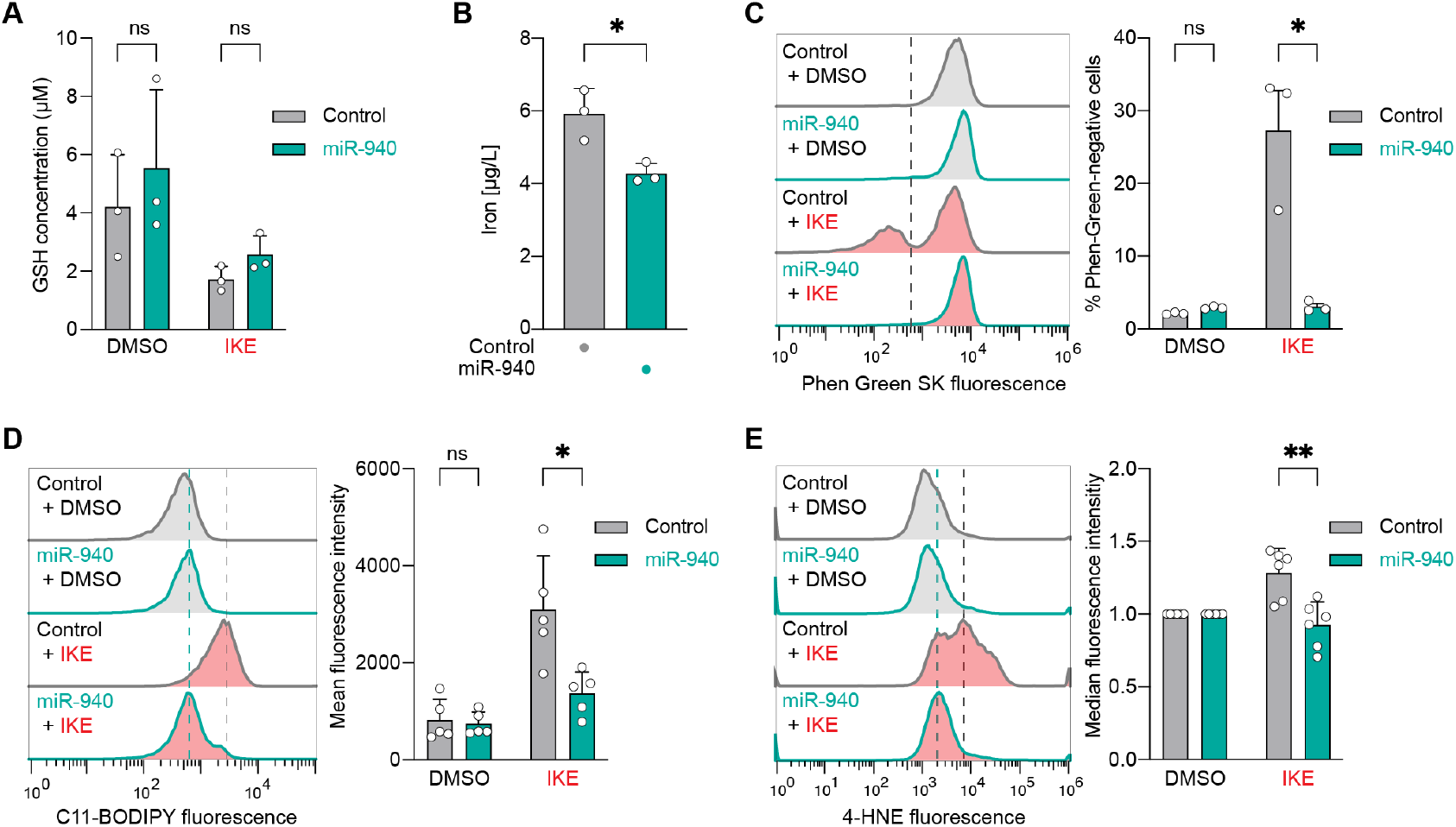
miR-940 suppresses iron accumulation and lipid peroxidation. A) Measurements of GSH concentration upon treatment with 2 µM IKE in both control and pre-miR-940 transfected H1080 cells, ns = p > 0.05, 2way ANOVA, data are mean ± SD of n = 3 biological replicates. B) Measurements of iron concentration via ICP-OES show reduced levels in miR-940-overexpressing cells compared to control cells, * = p ≤ 0.05, unpaired t-test with Welch’s correction, data are mean ± SD of n = 3 biological replicates. C) Analysis of iron levels via the heavy metal indicator PhenGreen SK Diacetate in pre-miR-940 transfected cells after treatment with 2 µM IKE compared to control cells, ns = p > 0.05, * = p ≤ 0.05, 2way ANOVA, data are mean ± SD of n = 3 biological replicates. D) C11-BODIPY sensor of lipid peroxidation upon treatment with 2 µM IKE in miR-940 overexpressing cells, ns = p > 0.05, * = p ≤ 0.05, 2way ANOVA, data are mean ± SD of n = 5 biological replicates. E) Detection of lipid peroxidation via antibody for 4-HNE in miR-940 overexpressing cells upon treatment with 2 µM IKE, ** = p ≤ 0.01, 2way ANOVA, data are mean ± SD of n = 6 biological replicates.

Next, iron levels of HT-1080 cells transfected with miR-940 were measured via ‘inductively coupled plasma optical emission spectroscopy’ (ICP-OES), as well as using the heavy metal indicator PhenGreen SK Diacetate. Both assays revealed a significant decrease in iron levels in cells overexpressing miR-940 before (Figure 4B) and after ferroptosis induction (Figure 4C). Moreover, lipid peroxidation was detected via staining with the C11-BODIPY sensor and a 4-HNE immunostaining after inducing ferroptotic cell death with IKE in HT-1080 cells. Both assays showed an induction of lipid peroxidation in IKE-treated control cells. Importantly, overexpression of miR-940 significantly suppressed levels of lipid peroxidation as detected by both techniques (Figure 4D, E). We also assessed lipid peroxidation using the C11-BODIPY sensor in HepG2 cells. These cells are more resistant to ferroptosis, which results in a smaller fluorescence shift. Nevertheless, a significant reduction of IKE-induced lipid peroxidation was detected in miR-940-overexpressing cells (Supplementary Figure 5A). Interestingly, similar to viability measurement (Figure 2B, 2F), miR-940 was not able to reduce lipid peroxidation induced by GPX4 inhibition (RSL3) in HT-1080 and HepG2 cells (Supplementary Figure 5B).

Together, these findings suggest that miR-940 inhibits ferroptotic cell death through three mechanisms: (1) enhanced repair of PLOOH due to upregulation of GPX4, (2) decreased levels of redox-active iron associated with reduced expression of NCOA4 and DMT1, and (3) a reduction in PUFA-PL levels through downregulation of LPCAT3 and ACSL4 (Figure 5).

## 3. Discussion

Our data suggest that miR-940 is a previously unrecognized cellular inhibitor of ferroptosis (Figure 2), acting through the coordinated modulation of the gene expression of ferroptosis regulators, especially the upregulation of the central gatekeeper of ferroptosis, namely GPX4. Rather than affecting individual effectors, it appears that miR-940 influences a broader regulatory architecture, the major hallmarks of ferroptosis, which controls cellular susceptibility to oxidative cell death. Therefore, we propose that miR-940 is a potent, novel cellular suppressor of ferroptotic cell death (Figure 5).

**Figure 5.**
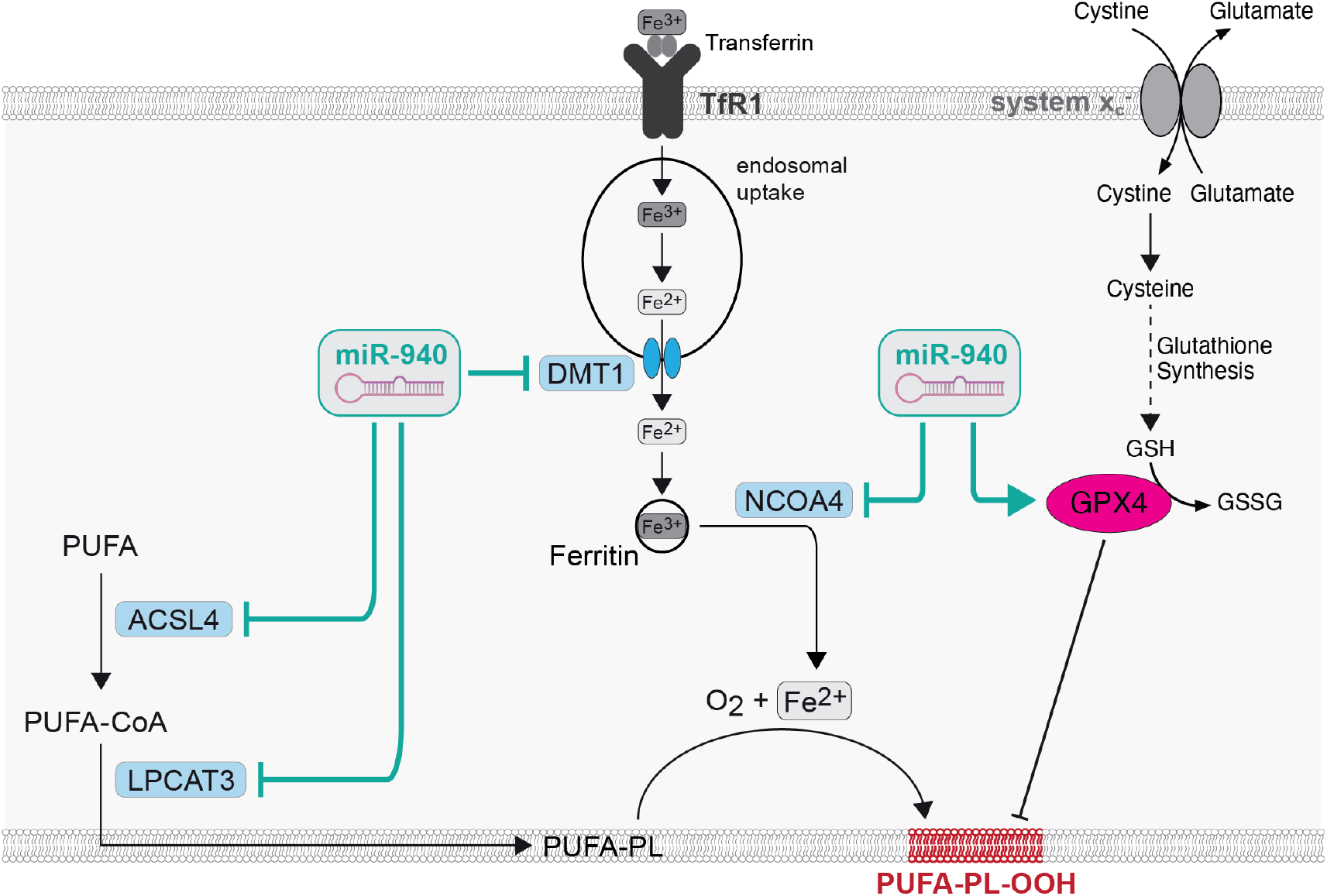
Working model of how miR-940 suppresses ferroptosis. This illustration shows how miR-940 suppresses ferroptosis by downregulating ACSL4 and LPCAT3, which leads to reduced PUFA-PL levels in the cell membrane. It also inhibits DMT1 to limit iron uptake, and NCOA4 to reduce ferritin degradation and thus the release of the labile, redox-active iron pool. This results in lower Fe^2^+ levels. Finally, it upregulates GPX4, which reduces lipid peroxides and counteracts ferroptosis.

Through integrative analysis, we mechanistically unravelled that miR-940 alters the expression profiles of several regulators involved in phospholipid processing and iron handling, both of which essentially govern vulnerability to lipid peroxidation (Figure 3). This suggests that small non-coding RNAs can have wide-ranging effects on cell fate by simultaneously targeting functionally linked molecular processes. Notably, ACSL4 and LPCAT3 are pivotal enzymes in determining phospholipid composition and susceptibility to peroxidation [3, 40], while DMT1 and NCOA4 influence intracellular iron availability [4], two critical nodes in ferroptotic sensitivity. The simultaneous repression of these targets by miR-940 suggests the existence of a robust evolutionarily mechanism to avoid lipid peroxidation driven cell death. Especially, the reduction of ACSL4 and LPCAT3 by miR-940 has the consequence that ferroptosis-relevant PUFA-containing phospholipids are largely depleted upon miR-940 overexpression as evident from lipidomics analysis (Figure 3F). Furthermore, the simultaneous upregulation of the selenoenzyme GPX4 by miR-940 (Figure 3) is interesting, given its central role in ferroptosis inhibition [1, 3-4]. Since miR-940 largely failed to protect cells from ferroptosis after GPX4 inhibition (viability and lipid peroxidation), these results suggest that GPX4 is the main axis through which miR-940 suppresses ferroptosis. The elevated levels of GPX4 protein upon miR-940 overexpression will still be susceptible to inhibition by RSL3, ML162, and FINO_2_. However, the exact mechanism by which miR-940 increases GPX4 expression remains unclear, it suggests the existence of indirect regulatory networks or the possibility of miR-940-mediated suppression of cellular GPX4 inhibitors; for example, an E3 ligase that triggers degradation of GPX4 by the proteasome. In some cases, miRNAs may also increase protein levels, which would be a direct effect. Thus, future investigation into upstream regulators and feedback loops involving miR-940 and GPX4 will help to decipher this process.

While this study provides mechanistic insight into the regulation of ferroptosis by miR-940, some limitations should be acknowledged. Our findings were confirmed in three independent cell lines, largely relying on *in vitro* experimental systems. To provide pathological context, we show that increased miR-940 expression correlates with decreased overall survival in several cancer patient cohorts, supporting the notion that inhibition of ferroptosis may negatively influence patient outcomes. However, direct functional validation *in vivo* is lacking. Future studies in animal models or patient-derived systems will assess the (patho)physiological significance of miR-940-mediated ferroptosis regulation. Lastly, although our data identify key factors associated with ferroptosis downstream of miR-940 and demonstrate that miR-940 does not influence apoptosis under our experimental conditions, potential off-target effects or additional roles of miR-940 in other cell death pathways cannot be completely ruled out.

From a translational perspective, miR-940 may represent a promising target in cancer entities such as sarcoma, liver hepatocellular carcinoma or kidney cancer (Figure 2K-M), where resistance to ferroptosis contributes to therapeutic failure. Inhibiting miR-940 may increase tumor sensitivity to ferroptosis inducers. In line with this, the elevated expression of miR-940 could serve as a predictive biomarker for identifying patients who are likely to respond to miR-940-modulating therapies in combination with ferroptosis-based medicines [41]. Combining miR-940 antagonism with ferroptosis-inducing regimens could increase anti-tumor efficacy and open new ways of overcoming drug resistance. These concepts provide a framework for future preclinical and clinical oncology studies exploring miR-940-targeted strategies.

In summary, this study identifies miR-940 as a potent regulator that limits ferroptotic cell death by reshaping lipid and iron homeostasis. These findings broaden our understanding of the role of microRNAs in controlling redox-sensitive death pathways and emphasize the therapeutic potential of targeting miR-940 in diseases associated with ferroptosis. Future efforts should focus on defining the upstream signals that govern miR-940 expression and analysing its interactions with other non-coding RNAs and transcription factors. A better understanding of this regulatory landscape will provide a deeper insight into the biology of ferroptosis and could reveal new opportunities for precision therapies in cancer.

## 4. Experimental Section

### Cell culture

HT-1080 cells (RRID:CVCL_0317), HepG2 cells (RRID:CVCL_0027) and HEK293T cells (RRID:CVCL_0063) were purchased from ATCC and grown in Dulbecco’s Modified Eagle’s medium (Thermo Fisher Scientific) containing 10% fetal bovine serum (Gibco), 1% Penicillin-Streptomycin (Thermo Fisher Scientific) and supplemented with 1% non-essential amino acids (Sigma) at 37°C with 5% CO_2_. Cells were regularly tested (including mycoplasma) and were completely contamination-free.

### Transfection of HT-1080 cells

If not indicated otherwise, 200,000 cells per well were seeded in 6-well plates the day before transfection. Cells were transfected with 30 nM hsa-miR-940 pre-miR or pre-miR negative control #1 (AM17100, Ambion, Invitrogen) in 1 mL medium. Per well, 3 μL of 10 μM miRNA or negative control were diluted in 50 μL Opti-MEM and 50 μL 6% Lipofectamine RNAiMAX reagent in Opti-MEM I reduced serum medium (all from Thermo Fisher Scientific) was added. The mix was incubated for 10 min at room temperature before dropwise addition to the cells. After another 5 min of incubation at room temperature, the plate was kept at 37 °C. After 6 h, cells were washed with PBS and fresh medium was added.

### Cell viability assays

Transfected cells were harvested 48 h after transfection with 0.05% Trypsin-EDTA and seeded into 384-well plates at a density of 750 cells per well. After incubation for 6 h, different concentrations of ferroptosis inducers (IKE (Cayman Chemical), RSL3 (Sigma Aldrich), ML162 (Sigma Aldrich), FinO_2_ (Caymen Chemical) or the apoptosis inducer staurosporine (Sigma Aldrich) were added to the cells. DMSO was used as a negative control. After 18 h at 37°C, CellTiter-Glo 2.0 Reagent (Promega) was added directly into each well and viability was assessed by luminescence measurement in an EnVision 2104 Multilabel plate reader (PerkinElmer).

### AnnexinV + 7AAD Detection Assay

HT-1080 cells were seeded in 6-well plates at a density of 100.000 cells per well one day prior to miRNA transfection, with transfections performed as previously described. In +STA conditions, 20 hours after transfection, 1 µM staurosporine was added to cultures for 4 hours. 24 hours after transfection, media was removed from all wells before being washed with 1x PBS and then incubated in 1 mL Accutase (Gibco) for 5 minutes at 37^°^C. Lifted cells were then collected in media and centrifuged at 300 x g for 3 minutes, washed with 1x PBS and again centrifuged at 300 x g for 3 minutes. AnnexinV assays were performed using the FITC Annexin V Apoptosis Detection Kit with 7-AAD (Biolegend, Cat # 640922), according to the kit protocol. Cell pellets were resuspended in 100 µL AnnexinV binding buffer along with 5 µL of AnnexinV and 5 µL of 7AAD, before being briefly vortexed and incubated in the dark at room temperature for 15 minutes. 400 µL of AnnexinV binding buffer was subsequently added to each tube, and samples were analyzed using the CytoFLEX S flow cytometer (Beckman Coulter), using the 488 nm laser with 525/40 bandpass filter to detect AnnexinV-FITC and the 561 nm laser with 610/20 bandpass filter to detect 7AAD. Data was analyzed using FlowJo (v10.10.0, BD Biosciences).

### Crystal Violet Staining

HT-1080 cells were seeded into a 6-well plate at a density of 200,000 cells per well and transfected as described. After incubation for 24 hours cells were treated with 2 µM IKE. DMSO was used as a control treatment. After 6 hours, the medium was removed, cells were washed with MilliQ water and 1.5 mL crystal violet staining solution (0.5% in 20% methanol) was added. Cells were incubated at room temperature for 20 min and rinsed with MilliQ water until no residual crystal violet solution was visible. Images were taken with an EVOS FL fluorescence microscope.

### Spheroid formation

Transfected HT-1080 cells were harvested 48 h after transfection and 3,000 cells per well were seeded into 96-well Round Bottom Ultra Low Attachment Microplates (Corning costar 7007). After 1 h of incubation at 37°C cells were treated with 2 µM IKE or DMSO as control. 48 h after treatment nuclei were stained with Hoechst 33342 (Sigma Aldrich) at a dilution of 1:10,000 followed by 1 h incubation at 37°C. Spheroids were imaged using the Operetta High Content Screening System (Perkin Elmer) at 10x magnification. Image analysis was performed with Columbus Software (Perkin Elmer).

### qRT-PCR

Total RNA was isolated from transfected cells 48 h after transfection using InVitrap Spin Universal RNA Mini kit (Stratec Scientific) according to the manufacturer’s instructions. Genomic DNA was removed by digestion with dsDNase (Promega) before reverse transcription of mRNA and miRNA. Maxima H Minus First Strand cDNA Synthesis Kit (Thermo Fisher Scientific) was used for synthesis of first-strand cDNA with oligo (dT)_18_ and random hexamer primers. Reverse transcription of miRNA was performed using MystiCq microRNA cDNA Synthesis Mix according to the manufacturer’s instructions. Polyadenylation of microRNA was performed with the Poly(A) Polymerase (Thermo Fisher Scientific, AM2030). Subsequently, RNA was eliminated by digestion with RNase H for 30 min at 37C°. qRT-PCR was performed on a LightCycler480 (Roche) with PowerUp SYBR Green Master Mix (Thermo Fisher Scientific) for cDNA transcribed from mRNA and SYBR Green Master Mix (Roche) for cDNA transcribed from miRNA. The primers used are listed in Supplementary Table 2. Expression levels of target genes were normalized to RNA Polymerase II and quantification was carried out using the delta delta Cp method. For expression levels of miRNA, SNORD44 was used for normalization, respectively.

### Western blotting

Transfected cells were harvested 48 h after transfection as described above, washed once with PBS and resuspended in 2x RotiLoad (Carl Roth) for lysis. The samples were sonified and proteins were denatured at 95°C for 5 min. After separation on 12.5% SDS-PAGE gels, proteins were transferred to PVDF membranes. Unspecific binding sites were blocked by incubation in 5% milk in TBS-Tween for 30 min at room temperature followed by incubation in primary antibody diluted in 2.5% milk in TBS-Tween at 4 °C overnight. Primary antibodies were ACSL4 (Santa Cruz, sc-365230, 1:200), DMT1 (Abcam, ab55735, 1:4000), GPX4 (Abcam, ab125066, 1:1000), NCOA4 (Abcam, ab86707, 1:2000), and β-Actin (Santa Cruz Biotechnology, sc-47778, 1:400). Membranes were washed three times for 10 min before incubation in secondary antibody (dilution 1:7,500 in milk-TBS-Tween) for 1 h at room temperature. Western Lightning ECL reagent (Perkin Elmer) was used for chemiluminescence detection. Densiometric analysis of protein bands was performed with ImageJ software.

Validation statements, relevant citations and antibody profiles are provided online by Santa Cruz Biotechnology and Abcam for each respective antibody used in this study.

### Detection of iron levels via PhenGreen

Transfected cells were harvested 48 h after transfection as described above and seeded in 6-well plates at a density of 500,000 cells per well. After 8 h incubation at 37°C ferroptosis was induced by 1.5 µM IKE for 17-18.5 h. Cells were incubated with 5 µM PhenGreen SK Diacetate (Thermo Fisher Scientific) in PBS for 10 min at 37°C in the dark. For dead cell exclusion 1 µg/mL propidium iodide (PI) was added to the cells 5 min before analysis on an Attune acoustic flow cytometer (Applied Biosystems). 15,000 PI-negative events were recorded, and analysis was performed with FlowJo v10.8.1 Software (BD Life Sciences).

### Detection of iron levels via ICP-OES

Cells were seeded at a density of 125,000 cells per 10 cm dish and transfected as described above. After 24 h incubation at 37°C ferroptosis was induced by 2 µM IKE for 18 h. Cells were harvested and centrifuged at 300 x g for 5 minutes. The cell pellet was washed with 5 mL PBS twice. The cell count was counted via Vi-CELL XR (Beckman Coulter) and adjusted to 1 Mio cells per sample in 1.5 mL ddH_2_O. Afterwards, 1.5 mL of 65% nitric acid were added to the samples and incubated at room temperature overnight. Samples were stored at 4°C. Iron levels were measured via inductively coupled plasma optical emission spectrometry (ICP-OES) using the Spectro Arcos 2 (Spectro Ametek).

### Detection of lipid peroxidation

Cells were prepared as described for detection of iron levels. However, for the analysis of lipid peroxidation 0.75 µM IKE was used with subsequent incubation at 37°C for 16.5-19 h. Afterwards BODIPY 581/591 C11 was added at a concentration of 2 µM followed by further 30 min incubation at 37°C in the dark. Dead cell exclusion and analysis were performed as described above.

For 4-HNE immunostaining transfected cells were treated with 2 µM IKE 30 h after transfection and incubated at 37°C for 16.5-19 h. Cells were harvested with 0.05% Trypsin-EDTA, washed with PBS and resuspended in 10 % normal goat serum (Thermo Fisher Scientific) for 30 min on ice. Cells were incubated with anti-4-HNE antibody (Abcam, ab46545, 1:50) for 1 h on ice, washed three times with PBS and incubated with anti-rabbit Alexa 488 antibody (Thermo Fisher Scientific, A32731, 1:200) for 30 min on ice. After washing twice with PBS, cells were resuspended in PBS and fluorescence was analyzed on an Attune acoustic flow cytometer (Applied Biosystems). Histograms and intensities were analyzed with FlowJo v10.8.1 Software (BD LifeSciences).

Validation statements, relevant citations and antibody profiles are provided online by Thermo Fisher Scientific and Abcam for each respective antibody used in this study.

### Detection of glutathione levels

HT-1080 cells were seeded in 6-well plates at a density of 150,000 cells per well and transfected as described above. After 48 h of incubation 3 µM IKE was applied to the cells for 6 h before harvest. Equal numbers of cells per condition were lysed in PBS containing 0.5% Nonidet-P40 and debris was removed by centrifugation (15 min, 20,000 x *g*, 4°C). The supernatant was transferred to a fresh tube and proteins were removed using Deproteinizing Sample Preparation Kit – TCA (Abcam) according to the manufacturer’s instructions. For the measurement of glutathione (GSH) levels the fluorimetric GSH/GSSG Ratio Detection Assay Kit II (Abcam) was used with black µClear 96-well plates with clear bottom. The preparation of standard curves and the assay procedure was performed according to the manufacturer’s instructions. GSH levels were obtained by fluorescence detection with an EnVision 2104 Multilabel plate reader (PerkinElmer).

### CRISPR knockout screen and data analysis

HT-1080-Cas9 cells were transduced with viruses containing the GeCKO v2 human CRISPR knockout pooled library [32] to generate the HT-1080-Cas9-gRNA-library cells. These cells were cultured in medium that was supplemented with 5 μg/mL of blasticidin and 0.7 μg/mL of puromycin. Next, the cells were expanded and on day-1, 18x T175 flasks were seeded at ∼20% confluency. By Day 3, when the cells reached ∼80% confluency, they were treated in duplicate as follows: DMSO (samples #1–2), IKE (0.8 μM, samples #3–4) and incubated for 20 h, then harvested on Day 4. The cells were washed once with PBS, detached using trypsin and the sample duplicates were pooled. A 300 µL aliquot of each sample was diluted 1:1 with medium for viability assessment (ViCell). The remaining cells were washed twice with PBS (with centrifugation steps) and flash-frozen on dry ice before storage at –80 °C and submitted to the MSKCC GES Core.

Genomic DNA (gDNA) was isolated from cell pellets using standard extraction protocols and the integrated guide RNA (gRNA) sequences were PCR-amplified from the extracted gDNA using primers that flanked the gRNA cassette. The amplicons were then purified, quantified and the resulting library was then subjected to next-generation sequencing on an Illumina HiSeq platform to determine gRNA representation across the two different conditions (DMSO versus IKE). Normalized guide frequencies amplified from viable cells were scored against control cells to identify enriched or depleted guides using ENCoRE [42]. In some instances, a full complement of guides for each transcript was not present in amplified samples and imputed values were utilized to calculate P-values, False Discovery Rate (FDR), and fold changes. Results from the CRISPR KO screen are in Supplementary Table 3.

### Analysis miRNA integration

To identify target genes of the enriched miRNAs, we queried both the miRTarBase [43] and TargetScan [44] databases using default parameters. Consecutively, the resulting 95 putative validated target genes from miRTarBase were exploited as seed genes in a protein network analysis, performed using the Weighted Protein-Protein Interaction (WPPI) Bioconductor package (Galhoz A, Turei D, P. Menden M (2022). wppi: Weighting protein-protein interactions. R package version 1.6.0, https://github.com/aGalhoz/wppi) with default settings (i.e., without disease-specific constraints besides the given seed genes). The top 10% highest scoring genes from WPPI were selected and integrated with the original miRNA target data from miRTarBase and TargetScan to expand the candidate gene list for downstream analyses.

### RNA sequencing analysis

RNA was isolated from transfected cells, 48 h after transfection with InVitrap Spin Universal RNA Mini kit (Stratec Scientific) according to the manufacturer’s instructions. 2.5 µg of each sample were shipped on dry ice to Genewiz (Azenta Life Sciences, Leipzig), where strand-specific RNA sequencing (Illumina Nova-Seq technology) with 20 million reads per sample and 2 x 150 bp read length was performed. Sample preparation included PolyA selection for rRNA removal. RNA sequencing data was subject to differential gene expression analysis leveraging the DESeq2 package and R software version 4.2.0. *P*-values were estimated using Wald test and adjusted with Benjamini-Hochberg correction. Associations were considered significant at False Discovery Rate (FDR) of 1% and absolute value of log2 fold-change (log2FC) > 1. The raw counts are in Supplementary Table 4 and the differential expression is shown in Supplementary Table 5.

### Proteomics analysis

Transfected cells were harvested 48 h after transfection and resuspended in 300 µL RIPA buffer containing 1 mM PMSF. After incubation on ice for 15 min the samples were sonified at 50% amplitude three times for 2 pulses. After another 15 min on ice lysates were centrifuged (5 min at 14,000 x *g*) and the amount of protein in the supernatants was determined by Pierce BCA Protein Assay Kit (Thermo Fisher Scientific).

10 µg of whole cell extract were digested using a modified filter aided sample preparation (FASP) digest procedure [45]. After reduction and alkylation using DTT and IAA, the proteins were centrifuged on a 30 kDa cutoff filter device (Sartorius), washed thrice with UA buffer (8 M urea in 0.1 M Tris/HCl pH 8.5) and twice with 50 mM ammoniumbicarbonate. The proteins were digested for 2 h at room temperature using 0.5 µg Lys-C (Wako Chemicals, Neuss, Germany) and for 16 h at 37°C using 1 µg trypsin (Promega). After centrifugation (10 min at 14,000 x *g*) the eluted peptides were acidified with 0.5% TFA and stored at −20°C.

LC-MS/MS analysis was performed on a Q-Exactive HF-X mass spectrometer (Thermo Fisher Scientific) online coupled to an Ultimate 3000 nano-RSLC (Thermo Fisher Scientific). Tryptic peptides were automatically loaded on a C18 trap column (300 µm inner diameter (ID) × 5 mm, Acclaim PepMap100 C18, 5 µm, 100 Å, LC Packings) at 30 µl/min flow rate prior to C18 reversed phase chromatography on the analytical column (nanoEase MZ HSS T3 Column, 100Å, 1.8 µm, 75 µm x 250 mm, Waters) at 250 nl/min flow rate in a 95 minutes non-linear acetonitrile gradient from 3 to 40% in 0.1% formic acid. Profile precursor spectra from 300 to 1500 m/z were recorded at 60000 resolution with an automatic gain control (AGC) target of 3e6 and a maximum injection time of 30 ms TOP15 fragment spectra of charges 2 to 7 were recorded at 15000 resolution with an AGC target of 1e5, a maximum injection time of 50 ms, an isolation window of 1.6 m/z, a normalized collision energy of 28 and a dynamic exclusion of 30 seconds.

Generated raw files were analyzed using MaxQuant 2.0.3.1. (MPI Biochemistry, Martinsried) with default settings [46], additionally including LFQ quantification with LFQ min. ratio count of 1 [47], quantification using unique peptides and application of the match-between-runs algorithm. Searches were performed within the MaxQuant software using the Andromeda search engine [48] against the Swissprot human protein database (20237 sequences, 11451954 residues) applying default search settings. Identifications were filtered for a peptide-spectrum-match and protein false discovery rate of each 1%. Missing values in the resulting list of identified protein groups were imputed from normal distribution separately for each intensity column in Perseus (MPI Biochemistry, Martinsried) [49]. Resulting protein LFQ intensities were used for calculation of fold-changes and statistical values applying a Student’s t-test.

### Lipidomics Analysis

Cells were seeded in 10-cm culture dishes at a density of 2 Mio cells per dish. After 24 h, cells were transfected with hsa-miR-940 pre-miR or pre-miR negative control as described previously. Cells were treated with DMSO or 2 µM IKE for 18 h. Each condition was performed in five biological replicates. For lipidomics sample preparation, cells were harvested on dry ice to minimize metabolic activity. Culture dishes were washed twice with 11 mL ice-cold PBS (dry ice), and all residual PBS was completely removed. Subsequently, 800 µL of ice-cold 80% methanol (HPLC grade), prepared in glass flasks and stored at −80 °C overnight, was added directly onto the cells and rapidly distributed across the dish. Cells were scraped using a rubber-tipped scraper, and the extract was transferred into collection tubes prepared on dry ice. Remaining cells were recovered with an additional 200 µL of 80% methanol and combined with the initial extract. Samples were stored at −80 °C. After thawing the cell samples, about 80 mg of glass beads were added to each sample to break down the cells and release cellular contents during homogenization. Homogenization was conducted using Preccellys24 (PeqLab) for three cycles of 20 seconds at 5500 rpm, with a 30-second break between cycles at 10 °C. 200 µL of cell homogenate was then transferred into 1.5 mL glass vials for further extraction. Three replicates of 20 µL homogenates were pipetted in a 96-well microtiter plate for Hoechst analysis [50].

Lipid extraction was performed by adding 60 µL water, 25 µL internal standard solution, and 575µL methyl tert-butyl ether (MTBE) to the sample homogenate in sequence, followed by incubation for 30 minutes on an orbital shaker DOS-10L (Neolabline, Heidelberg, Germany) at 300 rpm. For phase separation, 200 µL of water was added to each vial. The mixtures were vortexed, and the vials were centrifuged at 5,000 x g for 10 min at RT with a Sigma 4-5C centrifuge (Qiagen, Hilden, Germany). The upper (organic) phase was transferred into two 1.5 mL glass vials, with 250 µL in each vial. The organic solvent was then evaporated under nitrogen gas using a TurboVap® 96 dual evaporator (Biotage, Uppsala, Sweden). The aqueous phase was then reextracted by adding 100 µL methanol, 300 µL MTBE, and 100 µL water before the sample was incubated for 5 minutes at 300 rpm at room temperature. The sample was then centrifuged at 5,000 g for 10 minutes. The organic phase was transferred to the respective vials from the first extraction step (150 µL per vial) and evaporated to dryness under gaseous nitrogen. For quality control purposes, along with the biological samples, the pool of all homogenates, EDTA human reference plasma, and water were extracted using the same protocol as for the biological samples.

The dried extract samples were stored in −20°C until LC-MS analysis. Prior to LC-MS analysis, the dried extract was reconstituted in 40 µL of a mixture of 60% 2-propanol/ 35% acetonitrile/ 5% water. Lipid analysis was performed using a Sciex ExionAD LC system coupled to a Sciex ZenoTOF 7600 (Sciex, Darmstadt, Germany). Lipids were separated as described by Witting et al. [51]. Briefly, a Waters Cortecs UPLC C18 column (150 mm x 2.1 mm ID, 1.6 µm particle size) was used with eluent A being 40% H2O/60% ACN + 10 mM ammonium formate / 0.1% form acid and eluent B being 10% ACN/90%iPrOH + 10 mM ammonium formate / 0.1% formic acid. Samples have been analyzed in positive and negative ionization modes using data-dependent acquisition. Obtained raw data was processed using mzmine 4.8.30 [52], including conversion to .mzML format, peak picking, alignment and annotation. Lipids were annotated using rule-based lipid annotation in mzmine, and the results were exported as .csv files for further analysis in R and Excel. Data normalization based on Hoechst assay readouts and QC-based intensity drift correction was performed in a custom R workflow using the notame package [53]. After normalization, only features present in all QC samples with an RSD < 30% were kept for further statistical analysis. For statistical analysis Microsoft Excel (version 16.100) was used. The peak areas of the respective lipid features were log2 transformed and scaled relative to the mean values of all samples. A two-tailed t-test was performed to select significant (p ≤ 0.05) lipid changes and ferroptosis relevant lipids were chosen and presented in a heatmap created in the GraphPad Prism Software (version 10.6.1). Lipidomics data are in Supplementary Table 6.

## Statistical Analysis

Statistical analyses were performed using the GraphPad Prism Software (version 10.6.1). All relevant statistical information is provided in the respective figure legends.

## Acknowledgements

We thank Stefanie Brandner, Quirin Reinold and Norbert Fischer for excellent technical assistance. This work is funded by the Deutsche Forschungsgemeinschaft (DFG, German Research Foundation) – TRR 387/1 – 514894665 to K.H. This research was funded in part through the NIH/NCI Cancer Center Support Grant P30 CA0088748. This work was supported by NIH Shared Instrumentation Grants (S10OD012351, S10OD021764 and S10OD032433). Research reported in this publication was partially performed in the CCTI Flow Cytometry Core and the Molecular Pathology Shared Resource (MPSR) core, supported in part by the Office of the Director, National Institutes of Health under awards (S10OD020056). This research was also funded in part through the NIH/NCI Cancer Center Support Grant (P30CA013696) and used the Genomics and High Throughput Screening Shared Resource. This work was also funded in part through the NIH/NCI Cancer Center Support Grant (P30CA008748), the Columbia University Digestive and Liver Disease Research Center (funded by NIH grant 5P30DK132710) through use of its Bioimaging Core, Bioinformatic and Single Cell Analysis Core, Organoid and Cell Culture Core and Clinical Biospecimen and Research Core.

## Author Contributions

K.H. and B.R.S. initiated the study; K.H. oversaw and supervised the whole study; A.K., J.T., S.A.I.W., D.K., K.K., L.R., S.S. and K.H. performed experiments and analyzed the data; M.F. and R.G. performed NGS of gDNA from the CRISPR KO screen; J.A.S. analyzed the CRISPR KO screen; A.G. and M.P.M. performed bioinformatics analyses and analyzed the transcriptomics data; A.A. and M.W. performed lipidomics, and M.W. and J.T. analyzed the data; J.M-P. and S.M.H. performed proteomics and analyzed the data. M.W., H.Z., S.M.H., M.P.M., M.V., B.R.S., and K.H. supervised the work; S.A.I.W., A.K., and K.H. wrote the initial paper draft; All authors edited and finalized the paper.

## Data Availability Statement

The raw data of this study are available from the corresponding author upon reasonable request. All results from CRISPR KO screen are in Supplementary Table 3. The RNAseq data (count data) generated in this study are provided in the Supplementary Table 4 and 5. Lipidomics data are in Supplementary Table 6. Proteomics data are deposited at PRIDE database (Project accession: PXD064621).

## Code Availability

Source code can be downloaded from https://github.com/MendenLab/miRNAInt_Ferroptosis.

## Responsible use of Artificial Intelligence

Artificial intelligence (AI) tools including ChatGPT and DeepL were employed in this manuscript to refine grammar, spelling, and readability. These tools were used exclusively for linguistic enhancement, ensuring clarity and coherence without altering the scientific content, interpretation, or conclusions. All intellectual contributions, data analyses, and critical discussions remain the work of the authors. The final manuscript was thoroughly reviewed and approved by all co-authors to ensure accuracy and integrity.

## Legends for Supplementary Tables

**Supplementary Table 1**

Predicted miR-940 binding sites of the ferroptosis regulators NCOA4, ACSL4, LPCAT3, DMT1, and GPX4 by databases TargetScan and miRDB. Includes the site type, site count, site position, score and score percentile.

**Supplementary Table 2**

Table of primers used in qRT-PCR. Includes primers for expression of ACSL4, DMT1, GPX4, LPCAT3, miR-94 and RNA Pol II.

**Supplementary Table 3**

Differentially regulated miRNAs identified by the GeCKO v2 whole genome CRISPR knockout screen. Sheet 1 includes the whole data set of the CRISPR screen. Sheet 2 sows 15 miRNAs that were significantly enriched and 15 miRNAs that were significantly depleted.

**Supplementary Table 4**

Raw counts from strand-specific RNA sequencing of total RNA isolated from pre-miR-940- and control-transfected cells, generated using Illumina NovaSeq technology.

**Supplementary Table 5**

Differential expression analysis of the RNA-sequencing data. Analysis was performed using the DESeq2 package in R (version 4.2.0). P-values were estimated using the Wald test and adjusted for multiple testing using the Benjamini–Hochberg method. Associations were considered significant at a false discovery rate (FDR) of 1% and an absolute log2 fold change (|log2FC|) > 1.

**Supplementary Table 6**

Table of lipidomics data with statistical analysis in Microsoft Excel (version 16.100). The peak areas of the respective lipid features were log2 transformed and scaled relative to the mean values of all samples. A two-tailed t-test was performed to select significant (p ≤ 0.05) lipid changes.

## REFERENCES

[1] K. Hadian, B. R. Stockwell, Nat Rev Drug Discov 2023, 22 (9), 723.

[2] S. J. Dixon, K. M. Lemberg, M. R. Lamprecht, R. Skouta, E. M. Zaitsev, C. E. Gleason, D. N. Patel, A. J. Bauer, A. M. Cantley, W. S. Yang, B. Morrison, 3rd, B. R. Stockwell, Cell 2012, 149 (5), 1060.

[3] K. Hadian, B. R. Stockwell, Cell 2020, 181 (5), 1188.

[4] B. R. Stockwell, Cell 2022, 185 (14), 2401.

[5] J. Tschuck, V. Skafar, J. P. Friedmann Angeli, K. Hadian, Trends Biochem Sci 2025.

[6] C. Berndt, H. Alborzinia, V. S. Amen, S. Ayton, U. Barayeu, A. Bartelt, H. Bayir, C. M. Bebber, K. Birsoy, J. P. Böttcher, S. Brabletz, T. Brabletz, A. R. Brown, B. Brüne, G. Bulli, A. Bruneau, Q. Chen, G. M. DeNicola, T. P. Dick, A. Distéfano, S. J. Dixon, J. B. Engler, J. Esser-von Bieren, M. Fedorova, J. P. Friedmann Angeli, M. A. Friese, D. C. Fuhrmann, A. J. García-Sáez, K. Garbowicz, M. Götz, W. Gu, L. Hammerich, B. Hassannia, X. Jiang, A. Jeridi, Y. P. Kang, V. E. Kagan, D. B. Konrad, S. Kotschi, P. Lei, M. Le Tertre, S. Lev, D. Liang, A. Linkermann, C. Lohr, S. Lorenz, T. Luedde, A. Methner, B. Michalke, A. V. Milton, J. Min, E. Mishima, S. Müller, H. Motohashi, M. U. Muckenthaler, S. Murakami, J. A. Olzmann, G. Pagnussat, Z. Pan, T. Papagiannakopoulos, L. Pedrera Puentes, D. A. Pratt, B. Proneth, L. Ramsauer, R. Rodriguez, Y. Saito, F. Schmidt, C. Schmitt, A. Schulze, A. Schwab, A. Schwantes, M. Soula, B. Spitzlberger, B. R. Stockwell, L. Thewes, O. Thorn-Seshold, S. Toyokuni, W. Tonnus, A. Trumpp, P. Vandenabeele, T. Vanden Berghe, V. Venkataramani, F. C. E. Vogel, S. von Karstedt, F. Wang, F. Westermann, C. Wientjens, C. Wilhelm, M. Wölk, K. Wu, X. Yang, F. Yu, Y. Zou, M. Conrad, Redox Biol 2024, 75, 103211.

[7] H. Feng, K. Schorpp, J. Jin, C. E. Yozwiak, B. G. Hoffstrom, A. M. Decker, P. Rajbhandari, M. E. Stokes, H. G. Bender, J. M. Csuka, P. S. Upadhyayula, P. Canoll, K. Uchida, R. K. Soni, K. Hadian, B. R. Stockwell, Cell Rep 2020, 30 (10), 3411.

[8] J. Jin, K. Schorpp, D. Samaga, K. Unger, K. Hadian, B. R. Stockwell, ACS Chem Biol 2022, 17 (3), 654.

[9] K. Bersuker, J. M. Hendricks, Z. Li, L. Magtanong, B. Ford, P. H. Tang, M. A. Roberts, B. Tong, T. J. Maimone, R. Zoncu, M. C. Bassik, D. K. Nomura, S. J. Dixon, J. A. Olzmann, Nature 2019, 575 (7784), 688.

[10] S. Doll, F. P. Freitas, R. Shah, M. Aldrovandi, M. C. da Silva, I. Ingold, A. Goya Grocin, T. N. Xavier da Silva, E. Panzilius, C. H. Scheel, A. Mourão, K. Buday, M. Sato, J. Wanninger, T. Vignane, V. Mohana, M. Rehberg, A. Flatley, A. Schepers, A. Kurz, D. White, M. Sauer, M. Sattler, E. W. Tate, W. Schmitz, A. Schulze, V. O’Donnell, B. Proneth, G. M. Popowicz, D. A. Pratt, J. P. F. Angeli, M. Conrad, Nature 2019, 575 (7784), 693.

[11] E. Mishima, J. Ito, Z. Wu, T. Nakamura, A. Wahida, S. Doll, W. Tonnus, P. Nepachalovich, E. Eggenhofer, M. Aldrovandi, B. Henkelmann, K. I. Yamada, J. Wanninger, O. Zilka, E. Sato, R. Feederle, D. Hass, A. Maida, A. S. D. Mourão, A. Linkermann, E. K. Geissler, K. Nakagawa, T. Abe, M. Fedorova, B. Proneth, D. A. Pratt, M. Conrad, Nature 2022, 608 (7924), 778.

[12] V. A. N. Kraft, C. T. Bezjian, S. Pfeiffer, L. Ringelstetter, C. Müller, F. Zandkarimi, J. Merl-Pham, X. Bao, N. Anastasov, J. Kössl, S. Brandner, J. D. Daniels, P. Schmitt-Kopplin, S. M. Hauck, B. R. Stockwell, K. Hadian, J. A. Schick, ACS Cent Sci 2020, 6 (1), 41.

[13] M. Soula, R. A. Weber, O. Zilka, H. Alwaseem, K. La, F. Yen, H. Molina, J. Garcia-Bermudez, D. A. Pratt, K. Birsoy, Nat Chem Biol 2020, 16 (12), 1351.

[14] J. Tschuck, W. Tonnus, S. Gavali, A. Kolak, M. Mallais, F. Maremonti, M. Sato, I. Rothenaigner, J. P. Friedmann Angeli, D. A. Pratt, A. Linkermann, K. Hadian, Cell Death Dis 2024, 15 (11), 853.

[15] J. Tschuck, L. Theilacker, I. Rothenaigner, S. A. I. Weiß, B. Akdogan, V. T. Lam, C. Müller, R. Graf, S. Brandner, C. Pütz, T. Rieder, P. Schmitt-Kopplin, M. Vincendeau, H. Zischka, K. Schorpp, K. Hadian, Nat Commun 2023, 14 (1), 6908.

[16] J. Tschuck, V. Padmanabhan Nair, A. Galhoz, C. Zaratiegui, H. M. Tai, G. Ciceri, I. Rothenaigner, J. Tchieu, B. R. Stockwell, L. Studer, D. S. Cabianca, M. P. Menden, M. Vincendeau, K. Hadian, Nat Commun 2024, 15 (1), 7611.

[17] D. Liang, Y. Feng, F. Zandkarimi, H. Wang, Z. Zhang, J. Kim, Y. Cai, W. Gu, B. R. Stockwell, X. Jiang, Cell 2023, 186 (13), 2748.

[18] W. Tonnus, F. Maremonti, S. Gavali, M. N. Schlecht, F. Gembardt, A. Belavgeni, N. Leinung, K. Flade, N. Bethe, S. Traikov, A. Haag, D. Schilling, S. Penkov, M. Mallais, C. Gaillet, C. Meyer, M. Katebi, A. Ray, L. M. S. Gerhardt, A. Brucker, J. N. Becker, M. Tmava, L. Schlicker, A. Schulze, N. Himmerkus, A. Shevchenko, M. Peitzsch, U. Barayeu, S. Nasi, J. Putz, K. S. Korach, J. Neugarten, L. Golestaneh, C. Hugo, J. U. Becker, J. M. Weinberg, S. Lorenz, B. Proneth, M. Conrad, E. Wolf, B. Plietker, R. Rodriguez, D. A. Pratt, T. P. Dick, M. Fedorova, S. R. Bornstein, A. Linkermann, Nature 2025, 645 (8082), 1011.

[19] A. M. Mohr, J. L. Mott, Semin Liver Dis 2015, 35 (1), 3.

[20] A. Balihodzic, F. Prinz, M. A. Dengler, G. A. Calin, P. J. Jost, M. Pichler, Cell Death Differ 2022, 29 (6), 1094.

[21] H. Li, Y. Li, D. Tian, J. Zhang, S. Duan, Mol Ther Nucleic Acids 2021, 25, 53.

[22] P. Li, Z. Xiao, J. Luo, Y. Zhang, L. Lin, J Cell Mol Med 2019, 23 (4), 2475.

[23] K. Jiang, T. Zhao, M. Shen, F. Zhang, S. Duan, Z. Lei, Y. Chen, J Cancer 2019, 10 (12), 2735.

[24] X. Liu, X. Ge, Z. Zhang, X. Zhang, J. Chang, Z. Wu, W. Tang, L. Gan, M. Sun, J. Li, Oncotarget 2015, 6 (28), 25418.

[25] K. Su, C. F. Wang, Y. Zhang, Y. J. Cai, Y. Y. Zhang, Q. Zhao, Int J Oncol 2017, 50 (4), 1211.

[26] J. Zhao, Y. Hou, C. Yin, J. Hu, T. Gao, X. Huang, X. Zhang, J. Xing, J. An, S. Wan, J. Li, Oncogene 2020, 39 (8), 1724.

[27] R. Xu, F. Zhou, T. Yu, G. Xu, J. Zhang, Y. Wang, L. Zhao, N. Liu, Am J Transl Res 2019, 11 (12), 7351.

[28] D. Sun, L. Chen, H. Lv, Y. Gao, X. Liu, X. Zhang, Onco Targets Ther 2020, 13, 1569.

[29] S. Wang, L. Xu, X. Che, C. Li, L. Xu, K. Hou, Y. Fan, T. Wen, X. Qu, Y. Liu, FEBS Lett 2018, 592 (4), 621.

[30] J. Ma, F. Sun, C. Li, Y. Zhang, W. Xiao, Z. Li, Q. Pan, H. Zeng, G. Xiao, K. Yao, A. Hong, J. An, Cell Death Dis 2014, 5 (8), e1377.

[31] T. Xu, K. Zhang, J. Shi, B. Huang, X. Wang, K. Qian, T. Ma, T. Qian, Z. Song, L. Li, Am J Cancer Res 2019, 9 (2), 250.

[32] O. Shalem, N. E. Sanjana, E. Hartenian, X. Shi, D. A. Scott, T. Mikkelson, D. Heckl, B. L. Ebert, D. E. Root, J. G. Doench, F. Zhang, Science 2014, 343 (6166), 84.

[33] Y. Zhang, H. Tan, J. D. Daniels, F. Zandkarimi, H. Liu, L. M. Brown, K. Uchida, O. A. O’Connor, B. R. Stockwell, Cell Chem Biol 2019, 26 (5), 623.

[34] H. Wang, T. Song, Y. Qiao, J. Sun, Exp Ther Med 2020, 19 (2), 833.

[35] C. Yu, C. Zhang, L. Zhang, J. Gu, L. Qin, Tohoku J Exp Med 2025.

[36] Y. Fan, X. Che, K. Hou, M. Zhang, T. Wen, X. Qu, Y. Liu, Exp Cell Res 2018, 373 (1-2), 180.

[37] B. Győrffy, Br J Pharmacol 2024, 181 (3), 362.

[38] Innovation (Camb) 2024, 5 (3), 100625.

[39] S. Kishore, A. R. Gruber, D. J. Jedlinski, A. P. Syed, H. Jorjani, M. Zavolan, Genome Biol 2013, 14 (5), R45.

[40] S. Doll, B. Proneth, Y. Y. Tyurina, E. Panzilius, S. Kobayashi, I. Ingold, M. Irmler, J. Beckers, M. Aichler, A. Walch, H. Prokisch, D. Trümbach, G. Mao, F. Qu, H. Bayir, J. Füllekrug, C. H. Scheel, W. Wurst, J. A. Schick, V. E. Kagan, J. P. Angeli, M. Conrad, Nat Chem Biol 2017, 13 (1), 91.

[41] K. Hadian, B. R. Stockwell, Nat Chem Biol 2021, 17 (11), 1113.

[42] D. Trümbach, S. Pfeiffer, M. Poppe, H. Scherb, S. Doll, W. Wurst, J. A. Schick, BMC Genomics 2017, 18 (1), 905.

[43] H. Y. Huang, Y. C. Lin, S. Cui, Y. Huang, Y. Tang, J. Xu, J. Bao, Y. Li, J. Wen, H. Zuo, W. Wang, J. Li, J. Ni, Y. Ruan, L. Li, Y. Chen, Y. Xie, Z. Zhu, X. Cai, X. Chen, L. Yao, Y. Chen, Y. Luo, S. LuXu, M. Luo, C. M. Chiu, K. Ma, L. Zhu, G. J. Cheng, C. Bai, Y. C. Chiang, L. Wang, F. Wei, T. Y. Lee, H. D. Huang, Nucleic Acids Res 2022, 50 (D1), D222.

[44] S. E. McGeary, K. S. Lin, C. Y. Shi, T. M. Pham, N. Bisaria, G. M. Kelley, D. P. Bartel, Science 2019, 366 (6472).

[45] J. R. Wiśniewski, A. Zougman, N. Nagaraj, M. Mann, Nat Methods 2009, 6 (5), 359.

[46] S. Tyanova, T. Temu, J. Cox, Nat Protoc 2016, 11 (12), 2301.

[47] J. Cox, M. Y. Hein, C. A. Luber, I. Paron, N. Nagaraj, M. Mann, Mol Cell Proteomics 2014, 13 (9), 2513.

[48] J. Cox, N. Neuhauser, A. Michalski, R. A. Scheltema, J. V. Olsen, M. Mann, J Proteome Res 2011, 10 (4), 1794.

[49] S. Tyanova, T. Temu, P. Sinitcyn, A. Carlson, M. Y. Hein, T. Geiger, M. Mann, J. Cox, Nat Methods 2016, 13 (9), 731.

[50] C. Muschet, G. Möller, C. Prehn, M. H. de Angelis, J. Adamski, J. Tokarz, Metabolomics 2016, 12 (10), 151.

[51] M. Witting, T. V. Maier, S. Garvis, P. Schmitt-Kopplin, J Chromatogr A 2014, 1359, 91.

[52] R. Schmid, S. Heuckeroth, A. Korf, A. Smirnov, O. Myers, T. S. Dyrlund, R. Bushuiev, K. J. Murray, N. Hoffmann, M. Lu, A. Sarvepalli, Z. Zhang, M. Fleischauer, K. Dührkop, M. Wesner, S. J. Hoogstra, E. Rudt, O. Mokshyna, C. Brungs, K. Ponomarov, L. Mutabdśija, T. Damiani, C. J. Pudney, M. Earll, P. O. Helmer, T. R. Fallon, T. Schulze, A. Rivas-Ubach, A. Bilbao, H. Richter, L. F. Nothias, M. Wang, M. Orešič, J. K. Weng, S. Böcker, A. Jeibmann, H. Hayen, U. Karst, P. C. Dorrestein, D. Petras, X. Du, T. Pluskal, Nat Biotechnol 2023, 41 (4), 447.

[53] A. Klåvus, M. Kokla, S. Noerman, V. M. Koistinen, M. Tuomainen, I. Zarei, T. Meuronen, M. R. Häkkinen, S. Rummukainen, A. Farizah Babu, T. Sallinen, O. Kärkkäinen, J. Paananen, D. Broadhurst, C. Brunius, K. Hanhineva, Metabolites 2020, 10 (4).

