## Supplementary Figures for "miR-940 suppresses ferroptosis by controlling expression of key regulatory genes"

**for**

#### Supplementary Figure 1

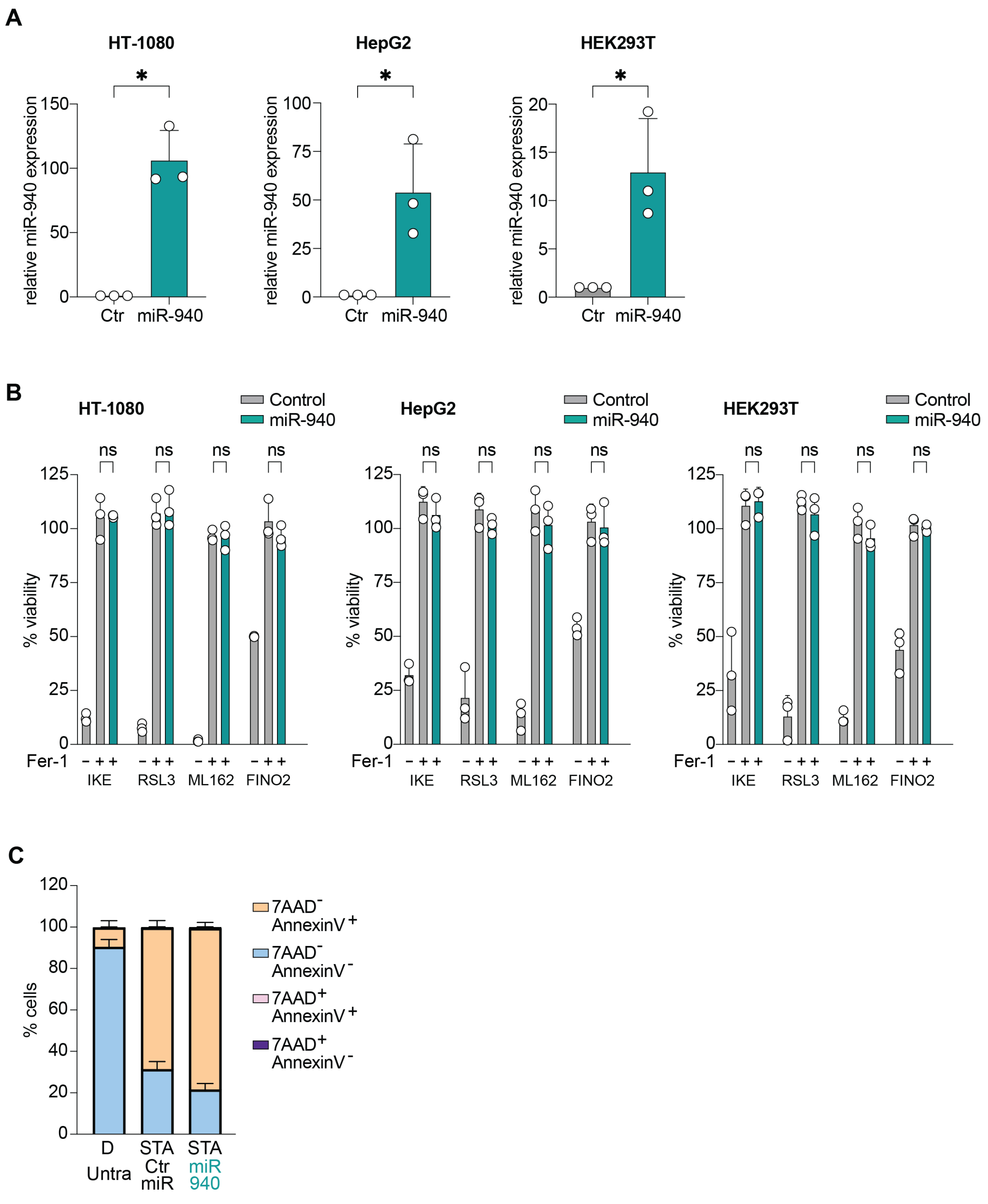


**Supplementary Figure 1: miR-940 inhibits IKE-induced ferroptosis**

**A)** Relative miR-940-expression in pre-miR-940- and control-transfected HT-1080, HepG2 and HEK293T cells, measured with qRT-PCR, * = $p \leq 0.05$, unpaired t-test with Welch's correction, data are mean ± SD of n = 3 biological replicates. **B)** Inhibition of ferroptosis in pre-miR-940 and control transfected HT-1080, HepG2, HEK293T treated with 0.3 µM IKE, 0.25 µM RSL3, 0.25 µM ML162 and 0.6 µM FINO2 (HT-1080), 2.5 µM IKE, 0.5 µM RSL3, 0.25 µM ML162 and 2,5 µM FINO2 (HepG2), 0.6 µM IKE, 0.25 µM RSL3, 0.25 µM ML162 and 1.25 µM FINO2 (HEK293T) with 2 µM Fer-1, ns = $p > 0.05$, 2way ANOVA, data are mean ± SD of n = 3 biological replicates. **C)** 7-AAD/AnnexinV-Staining in pre-miR-940- , control-transfected and untransfected HT-1080 cells 24h post transfection treated with Staurosporine (STA, 1 µM), data are mean ± SD of n = 3 biological replicates.

### Supplementary Figure 2

- **TargetScan**:
    - *NCOA4*: predicted pairing at position 117-124 of the 3'UTR (RNA sequence **CCUGCCUA**, hsa-miR-940 sequence **GGACGGAA**).

<table>
<tr><th></th><th>Predicted consequential pairing of target region (top) and miRNA (bottom)</th><th>Site type</th><th>Context++ score</th><th>Context++ score percentile</th><th>Weighted context++ score</th><th>Conserved branch length</th><th>$P_{CT}$</th><th>Predicted relative $K_D$</th></tr>
<tr><td>Position 117-124 of NCOA4 3' UTR<br>hsa-miR-6893-5p</td><td>5' ...UAGCUUAGUUCUUCUCCUGCCUA...<br>|||||||<br>3' CGAGGUGGGAUGUGGACGGAC</td><td>8mer</td><td>-0.37</td><td>99</td><td>-0.37</td><td>0.075</td><td>N/A</td><td>N/A</td></tr>
<tr><td>Position 117-124 of NCOA4 3' UTR<br>hsa-miR-6808-5p</td><td>5' ...UAGCUUAGUUCUUCUCCUGCCUA...<br>|||||||<br>3' GUACCAGGGUGGAGGGACGGAC</td><td>8mer</td><td>-0.37</td><td>99</td><td>-0.37</td><td>0.075</td><td>N/A</td><td>N/A</td></tr>
<tr><td>Position 117-124 of NCOA4 3' UTR<br>hsa-miR-940</td><td>5' ...UAGCUUAGUUCUUCUCCUGCCUA...<br>|||||||<br>3' CCCCUCGCCCCCGGGACGGAA</td><td>8mer</td><td>-0.37</td><td>99</td><td>-0.37</td><td>0.075</td><td>N/A</td><td>N/A</td></tr>
</table>

    - *ACSL4*: predicted pairing at position 2272-2278 of the 3'UTR (RNA sequence **CCUGCCU**, hsa-miR-940 sequence **GGACGGA**).

<table>
<tr><th></th><th>Predicted consequential pairing of target region (top) and miRNA (bottom)</th><th>Site type</th><th>Context++ score</th><th>Context++ score percentile</th><th>Weighted context++ score</th><th>Conserved branch length</th><th>$P_{CT}$</th><th>Predicted relative $K_D$</th></tr>
<tr><td>Position 2272-2278 of ACSL4 3' UTR<br>hsa-miR-6893-5p</td><td>5' ...AAUUCCAUGAAUGUUCCUGCCUC...<br>|||||||<br>3' CGAGGUGGGAUGUGGACGGAC</td><td>7mer-m8</td><td>-0.18</td><td>91</td><td>-0.10</td><td>0.043</td><td>N/A</td><td>N/A</td></tr>
<tr><td>Position 2272-2278 of ACSL4 3' UTR<br>hsa-miR-6808-5p</td><td>5' ...AAUUCCAUGAAUGUUCCUGCCUC...<br>|||||||<br>3' GUACCAGGGUGGAGGGACGGAC</td><td>7mer-m8</td><td>-0.17</td><td>89</td><td>-0.09</td><td>0.043</td><td>N/A</td><td>N/A</td></tr>
<tr><td>Position 2272-2278 of ACSL4 3' UTR<br>hsa-miR-940</td><td>5' ...AAUUCCAUGAAUGUUCCUGCCUC...<br>|||||||<br>3' CCCCUCGCCCCCGGGACGGAA</td><td>7mer-m8</td><td>-0.14</td><td>86</td><td>-0.08</td><td>0.043</td><td>N/A</td><td>N/A</td></tr>
</table>

    - *LPCAT3*: two predicted pairings at positions 273-279 and 650-656 of the 3'UTR (RNA sequence **CCUGCCU**, hsa-miR-940 sequence **GGACGGA**).

<table>
<tr><th></th><th>Predicted consequential pairing of target region (top) and miRNA (bottom)</th><th>Site type</th><th>Context++ score</th><th>Context++ score percentile</th><th>Weighted context++ score</th><th>Conserved branch length</th><th>$P_{CT}$</th><th>Predicted relative $K_D$</th></tr>
<tr><td>Position 273-279 of LPCAT3 3' UTR<br>hsa-miR-940</td><td>5' ...GGUUCAAGUGAUUUUCCUGCCUC...<br>|||||||<br>3' CCCCUCGCCCCCGGGACGGAA</td><td>7mer-m8</td><td>-0.20</td><td>94</td><td>-0.20</td><td>0.073</td><td>N/A</td><td>N/A</td></tr>
<tr><td>Position 273-279 of LPCAT3 3' UTR<br>hsa-miR-6893-5p</td><td>5' ...GGUUCAAGUGAUUUUCCUGCCUC...<br>|||||||<br>3' CGAGGUGGGAUGUGGACGGAC</td><td>7mer-m8</td><td>-0.20</td><td>93</td><td>-0.20</td><td>0.073</td><td>N/A</td><td>N/A</td></tr>
<tr><td>Position 273-279 of LPCAT3 3' UTR<br>hsa-miR-6808-5p</td><td>5' ...GGUUCAAGUGAUUUUCCUGCCUC...<br>|||||||<br>3' GUACCAGGGUGGAGGGACGGAC</td><td>7mer-m8</td><td>-0.19</td><td>92</td><td>-0.19</td><td>0.073</td><td>N/A</td><td>N/A</td></tr>
<tr><td>Position 650-656 of LPCAT3 3' UTR<br>hsa-miR-940</td><td>5' ...AGAGGGGUGCAAGGCCCUGCCUG...<br>|||||||<br>3' CCCCUCGCCCCCGGGACGGAA</td><td>7mer-m8</td><td>-0.05</td><td>66</td><td>-0.05</td><td>0.073</td><td>N/A</td><td>N/A</td></tr>
</table>

    - *DMT1*: predicted pairing at position 1433-1439 of the 3'UTR (RNA sequence **CCUGCCU**, hsa-miR-940 sequence **GGACGGA**).

<table>
<tr><th></th><th>Predicted consequential pairing of target region (top) and miRNA (bottom)</th><th>Site type</th><th>Context++ score</th><th>Context++ score percentile</th><th>Weighted context++ score</th><th>Conserved branch length</th><th>$P_{CT}$</th><th>Predicted relative $K_D$</th></tr>
<tr><td>Position 1433-1439 of SLC11A2 3' UTR<br>hsa-miR-940</td><td>5' ...ACCUCAAGUGAUCUACCUGCCUC...<br>|||||||<br>3' CCCCUCGCCCCCGGGACGGAA</td><td>7mer-m8</td><td>-0.12</td><td>83</td><td>0.00</td><td>0.099</td><td>N/A</td><td>N/A</td></tr>
</table>

    - *GPX4*: predicted pairing at position 98-104 of the 3'UTR (RNA sequence **CCUGCCU**, hsa-miR-940 sequence **GGACGGA**).

<table>
<tr><th></th><th>Predicted consequential pairing of target region (top) and miRNA (bottom)</th><th>Site type</th><th>Context++ score</th><th>Context++ score percentile</th><th>Weighted context++ score</th><th>Conserved branch length</th><th>$P_{CT}$</th><th>Predicted relative $K_D$</th></tr>
<tr><td>Position 98-104 of GPX4 3' UTR<br>hsa-miR-940</td><td>5' ...CGGCACUCAUGACGGCCUGCCUG...<br>|||||||<br>3' CCCCUCGCCCCCGGGACGGAA</td><td>7mer-m8</td><td>-0.25</td><td>97</td><td>-0.25</td><td>0.016</td><td>N/A</td><td>N/A</td></tr>
</table>

- **miRDB**: We identified one high-confidence site in *NCOA4* (8-mer, score = 84 out of 100), convergent with the TargetScan prediction at the same 3'UTR position (117-124).

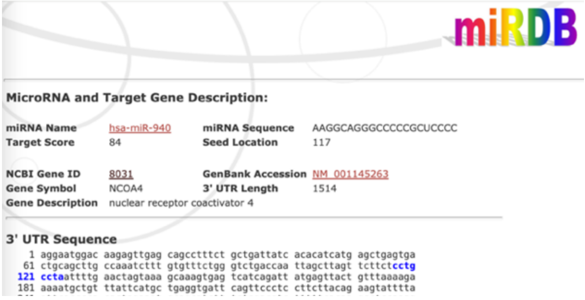


- **miRTarBase**: We identified a validated functional interaction between *DMT1* and miR-940 in the 3'UTR.

**Supplementary Figure 2: Prediction of direct binding sites for mature hsa-miR-940 sequence**
Prediction of direct binding sites for mature hsa-miR-940 sequence from miRbase as well as evaluated predicted binding sites using TargetScan (predicted targets), miRDB (predicted targets), and miRTarBase (validated targets). TargetScan predicted pairing of targets NCOA4, ACSL4, LPCAT3, DMT1 and GPX4. In addition, the miRDB identified the target NCOA4 and the miRTarBase the target DMT1.

#### Supplementary Figure 3

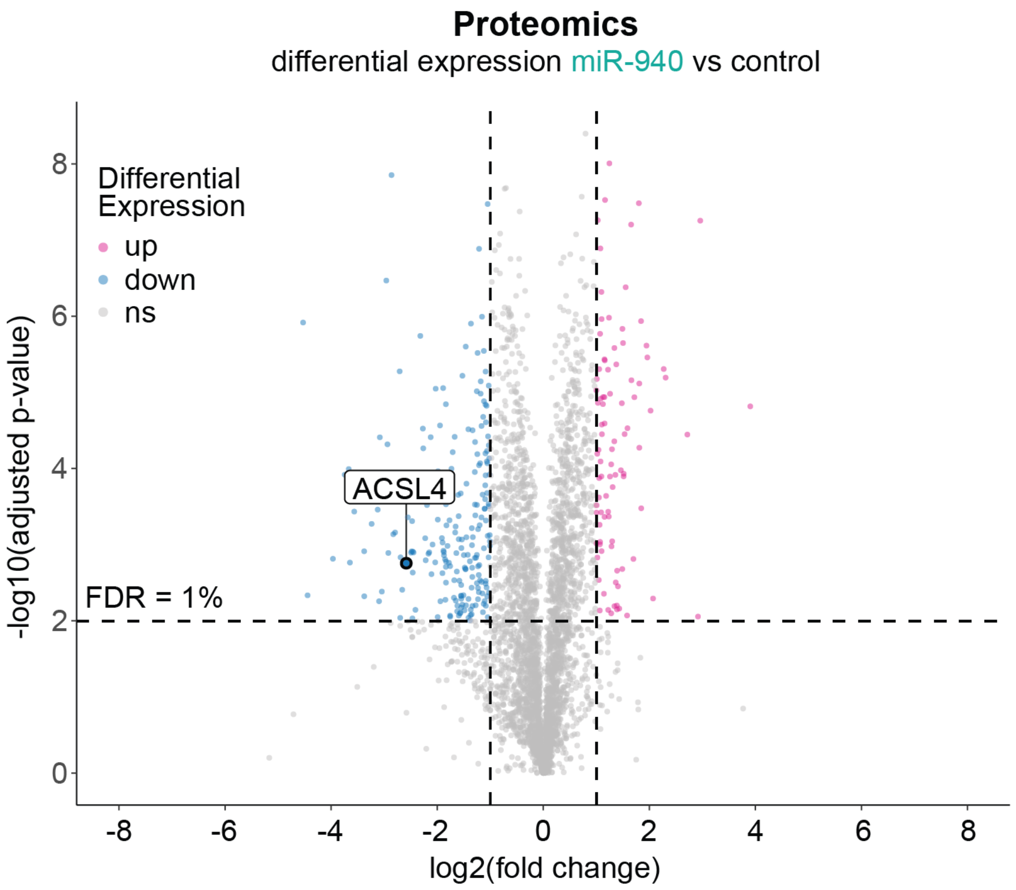


**Supplementary Figure 3: Proteomics analysis of miR-940 transfected cells**
Proteomics measurements of HT-1080 cells transfected with pre-miR-940 show downregulation of ACSL4 compared to control cells, FDR of 1%, Student's t-test.

### Supplementary Figure 4

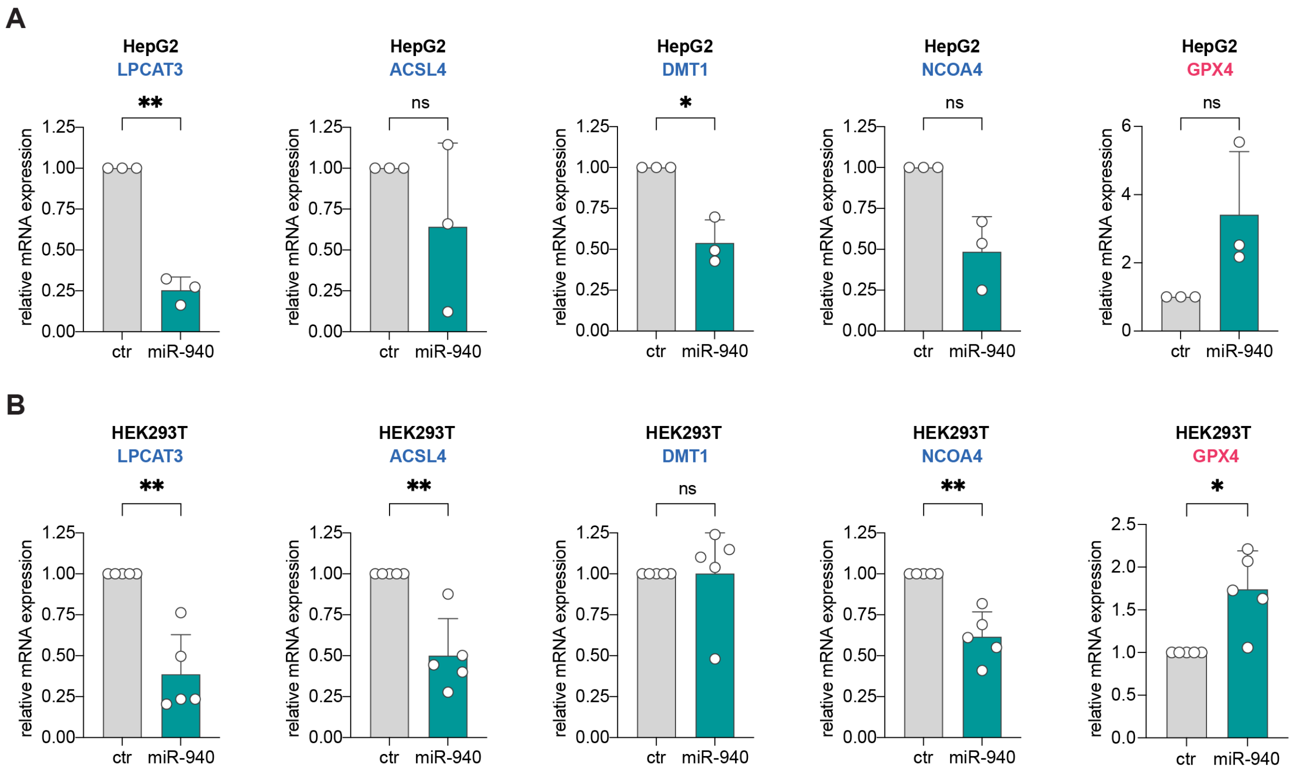


**Supplementary Figure 4: miR-940 modulates key regulators of ferroptotic cell death in HepG2 and HEK293T cells**
**A,B)** qRT-PCR of LPCAT3, ACSL4, DMT1, NCOA4, and GPX4 mRNA levels after pre-miR-940 transfection in HepG2 (A) and HEK293T (B) cells; ns = $p > 0.05$, * = $p \leq 0.05$, ** = $p \leq 0.01$ 2way ANOVA, data are mean ± SD of (A) n = 3 biological replicates and (B) 5 biological replicates.

#### Supplementary Figure 5

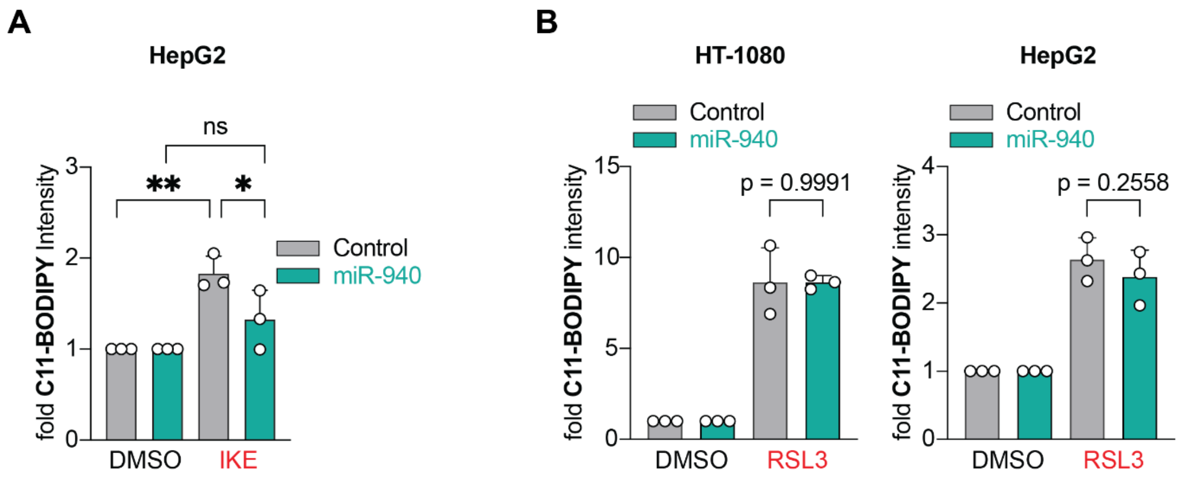


**Supplementary Figure 5: miR-940 suppresses lipid peroxidation**

**A)** C11-BODIPY sensor of lipid peroxidation upon treatment with 2 µM IKE in miR-940 overexpressing HepG2 cells, ns = $p > 0.05$, * = $p \leq 0.05$, ** = $p \leq 0.01$ 2way ANOVA, data are mean ± SD of n = 3 biological replicates. **B)** C11-BODIPY sensor of lipid peroxidation upon treatment with 150 nM RSL3 in miR-940 overexpressing HT-1080 cells as well as 1 µM RSL3 in miR-940 overexpressing HepG2 cells; 2way ANOVA, data are mean ± SD of n = 3 biological replicates.
